# Towards a universal spatial molecular atlas of the mouse brain

**DOI:** 10.1101/2024.05.27.594872

**Authors:** Yichun He, Hao Sheng, Hailing Shi, Wendy Xueyi Wang, Zefang Tang, Jia Liu, Xiao Wang

## Abstract

Spatially mapping a comprehensive brain at the single-cell level with molecular resolution is crucial for understanding its biological structure, mechanisms, and functions. Recent scalable spatial transcriptomics technologies^1-7^ have revolutionized our ability to examine millions of cells within the complex brain network, allowing for the generation of multiple mouse brain atlases that illuminate diverse transcriptomic and spatial cell types. However, despite enormous efforts, these spatial transcriptomics atlases still face challenges in providing a comprehensive brain representation: each atlas only captures a fraction of the vast cellular population of the brain and is limited in one or more aspects of spatial resolution, gene detection sensitivity, and gene throughput^8^. Here, we introduce FuseMap, a deep-learning-based framework for spatial transcriptomics that bridges single-cell or single-spot gene expression with spatial contexts and consolidates various gene panels across spatial transcriptomics atlases. By training on an extensive spatial transcriptomics atlas corpus of the mouse brain^1-7^, comprising over 18.6 million cells or spots and 26,665 genes from 434 tissue sections across seven datasets in a self-supervised way, FuseMap gains a fundamental understanding of cell identities and gene characteristics. We thus create a comprehensive molecular spatial reference atlas of the whole mouse brain with transcriptome-wide information, achieving multiple tasks including nomenclature harmonization of molecular cell types and tissue regions, identification of novel molecular brain regions, spatial gene imputation, targeted gene-panel selection, and region-specific cell-type interactions inference. Furthermore, the pretrained brain FuseMap model enables mapping new query data to the molecular spatial reference brain through transfer learning, allowing for automated cell and tissue annotations and replacing laborious and suboptimal manual annotations. Overall, we offer FuseMap as a novel computational framework for integration across various spatial transcriptomics platforms and provide a molecular common coordinate framework (molCCF) with pretrained FuseMap brain model for exploration of the brain molecular and cellular landscape with harmonized annotations and coordinates.

## Main text

Understanding the spatial complexity of the mammalian brain at molecular and cellular levels is fundamental for understanding the molecular architecture of its anatomy, function, and disorders. Advances in spatial transcriptomics technologies have transformed our ability to explore the molecular diversity of the mammalian brain, composed of millions to billions of cells. Recently, this transformation has facilitated the generation of multiple spatial atlases of the adult mouse brain, providing detailed insights into molecularly defined cell types and tissue regions across the three-dimensional (3D) brain^1-7^. However, each spatial transcriptomics atlas was built by a different technological platform that only captured a portion of the mouse brain. They also varied in spatial resolution, detection sensitivity, and gene panels^8^. As a result, building a common brain atlas that integrates all existing atlases is essential, from which we can gain a comprehensive view of the brain’s molecular architecture, cell types, and spatial organizations. Furthermore, as the generation of new large-scale spatial transcriptomics datasets becomes more routine, it is important to establish a unified reference atlas for researchers to navigate the brain^9^ and a pretrained model toward a foundation model to integrate, analyze, and interpret new data. Through transfer learning, such a foundation model will further enable automated annotations of new datasets, bypassing the labor-intensive, inaccurate, and inefficient process of manual data labeling^10,11^. Therefore, it is highly desired to create a universal reference of the mouse brain by integrating multiple spatial transcriptomics atlases from various technological platforms for large-scale integration and cross-atlas annotation.

However, it is challenging to use existing methodologies^12-25^ to integrate or annotate multiple atlas-scale, brain-wide spatial transcriptomics datasets. First, the primary challenge arises from the unique dual-modal nature of spatial transcriptomics data, which captures both the molecular signature of single-cell or single-spot gene expression profiles and their geospatial contexts through physical coordinates. Second, data integration of image-based spatial transcriptomics experiments is limited by the partial overlap in targeted gene panels across different atlases, leading to ambiguity in identifying cell types and tissue regions. Third, the nonlinear, technology-dependent, and noise-impacted expression variations further complicate the integration task. Thus, an accurate and effective spatial transcriptomics integration is required to preserve biological variation while correcting for batch effects across disparate spatial transcriptomics atlases^26^. In addition, it is also challenging to use existing methods to annotate spatial atlas from the reference map, as current automated cell annotation transfer methods are developed for single-cell RNA sequencing (scRNA-seq) data^10,11,27,28^ without incorporating spatial information. Altogether, there is an urgent need to build a scalable framework that can integrate large spatial transcriptomics atlases by leveraging both spatial and single-cell molecular characteristics. Furthermore, this framework can generate a pretrained model for automated integration and efficient annotation of new datasets through transferring the knowledge from reference atlases.

Here, we present a comprehensive brain representation by establishing a deep-learning framework, FuseMap, that overcomes these challenges and trains on multiple spatial transcriptomics mouse brain atlases (Fig. 1). FuseMap generates universal representations of genes, cells, and tissues with a self-supervised learning approach that eliminates the need for existing cell annotations. Specifically, it learns a unified embedding of cells in both single-cell transcriptomic profiles and spatial contexts, which are linked through physical tissue coordinates of cells; and it creates a universal gene embedding that consolidates all spatial transcriptomics gene panels (Fig. 1b,c). We trained FuseMap on seven publicly available spatial transcriptomics atlases of the mouse brain generated by different technologies^1-7^: Shi et al.^1^ (Atlas 1, STARmap PLUS), Yao et al.^2^ (Atlas 2, MERSCOPE), Zhang et al.^3^ (Atlas 3, MERFISH), Vizgen^4^ (Atlas 4, MERSCOPE), Borm et al.^5^ (Atlas 5, EEL-FISH), Chen et al.^6^ (Atlas 6, Stereo-seq), Langlieb et al.^7^ (Atlas 7, Slide-seq V2) (Fig. 1a, Supplementary Table 1). The pretrained FuseMap brain model gained a fundamental understanding of the spatial cellular organization of the whole mouse brain, providing universal embeddings of 18,637,640 cells/spots and 26,665 genes across 434 tissue sections. Based on the embeddings, we created a comprehensive molecular brain common coordinate framework (molCCF) (Fig. 1d). At the single-cell level, we harmonized cross-atlas molecular cell-type nomenclatures. At the spatial level, we unified and refined molecular brain regions marked by enriched cells and genes. At the gene level, we not only predicted transcriptome-wide gene expression but also revealed single-cell and spatially informed co-regulatory gene modules that enable informative targeted gene panel selection. Together with the harmonized cell types, tissue regions, and transcriptome-wide gene expression in the molCCF, we investigated distinct subtypes under different microenvironments, including spatially distinct cell subtypes of mature oligodendrocytes between white and grey matter. We also explored cell-cell interactions or communications, revealing consistent interaction patterns across different brain atlases. Furthermore, we demonstrated that the pretrained brain FuseMap model together with the molCCF allows users to annotate new query data with cell-type and brain-region labels, through accurate and efficient transfer learning (Fig. 1e).

**Figure 1.**
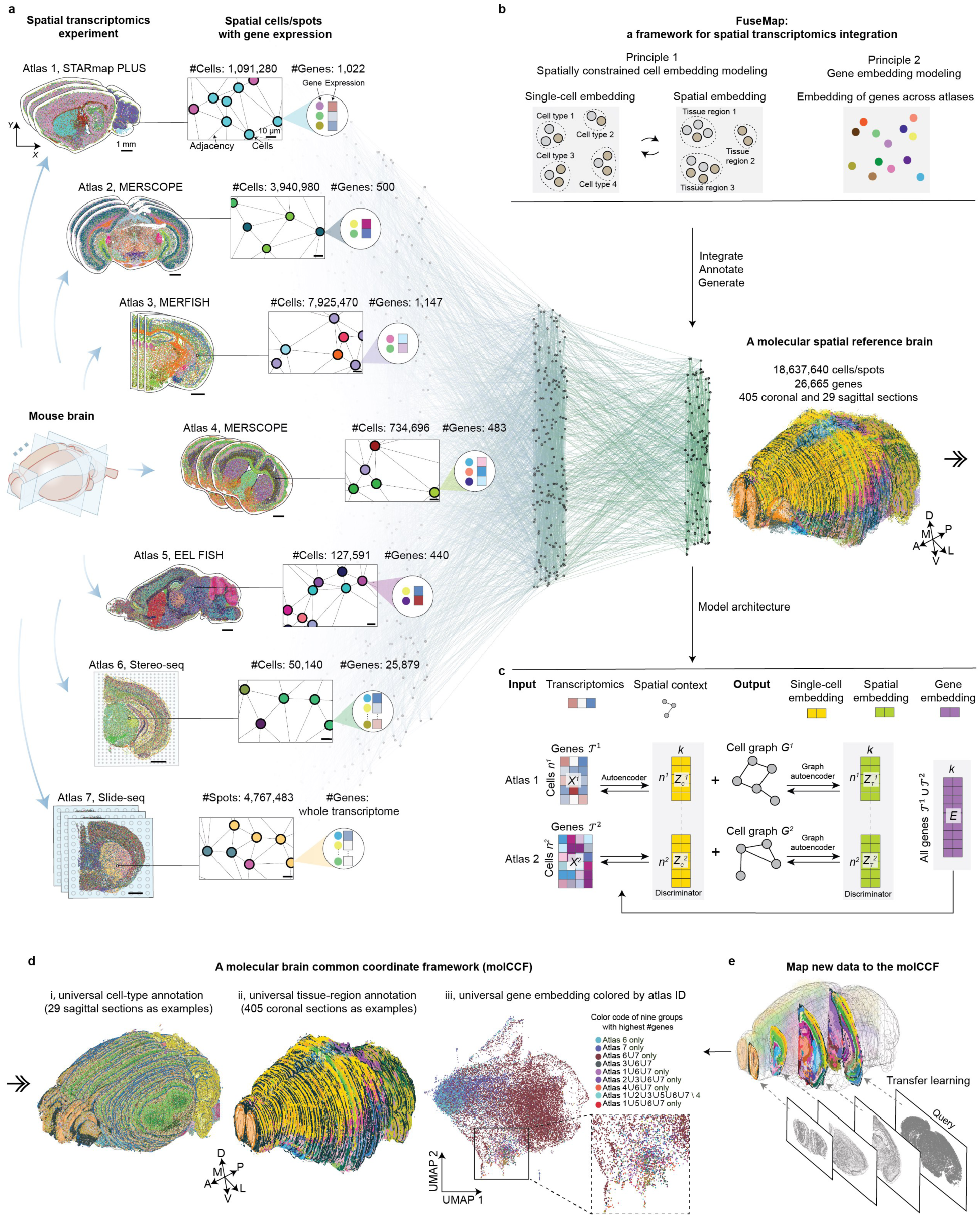
FuseMap as a spatial transcriptomics deep-learning model for atlas integration. **a**, Schematic showing that FuseMap integrates, annotates, and generates a comprehensive molecular spatial atlas corpus of over 18.6 million cells or spots and 26,665 genes across multiple spatial brain atlases. **b**, Key features in FuseMap designed to overcome challenges in spatial transcriptomics data integration. **c**, Schematic illustration of the model architecture of FuseMap. **d**, A molecular brain common coordinate framework (molCCF) with universal cell types, tissue regions, and gene embedding generated by FuseMap. **e**, FuseMap enabling the mapping of new data to the molCCF by transfer learning.

## Principles and model architecture of FuseMap

FuseMap addresses the challenges in spatial transcriptomics data integration in two aspects (Fig. 1b, Methods). First, it learns low-dimensional embeddings^1^ for each cell from the measured single-cell transcriptomic modality (termed as single-cell embedding) and spatially informed gene expression modality (termed as spatial embedding), which are paired via the unique physical tissue coordinates. This dual-modality modeling approach seamlessly links cellular gene expression with spatial contexts, facilitating a comprehensive characterization of cell identity. Second, FuseMap models gene characteristics as low-dimensional embeddings^29,30^ that inform the spatial expression patterns, and learns a universal gene embedding across various datasets. It leverages distinct genes from each dataset while connecting them through shared gene sets, thereby addressing the challenges of non-overlapping gene panels across spatial transcriptomics datasets.

Specifically, in the FuseMap framework (Fig. 1c and Methods), the transcriptomic data of each spatial transcriptomics sample, represented as a cell or spot by gene matrix 𝑋, is first encoded into a single-cell embedding 𝑍_𝑐_within a low-dimensional latent space. Then spatial locations of cells/spots construct a neighborhood graph 𝐺, connecting those in proximity^31^. The single-cell embedding 𝑍_𝑐_ and spatial graph 𝐺 are aggregated through a graph convolutional network to generate a spatial embedding 𝑍_𝑇_ within a spatially aware latent space. Conversely, 𝑍_𝑇_is converted back into 𝑍_𝑐_by decoupling the spatial graph 𝐺 through decoders. Finally, 𝑍_𝑐_ is transformed back into the original transcriptomic space, when combined with a learned gene embedding 𝐸, to reconstruct the gene expression matrix 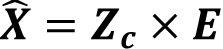, akin to coupled matrix factorization. The learning process entails adversarial optimization of autoencoder and discriminator losses: the autoencoder ensures that both transcriptomic information and spatial contexts are retained in 𝑍_𝑐_ and 𝑍_𝑇_, while the discriminator minimizes the distance between embeddings from different tissue samples for batch correction. Each tissue section’s unique characteristics were modeled using dedicated autoencoders and graph autoencoders, forming a section-specific probabilistic generative model. Collectively, FuseMap is trained in an entirely self-supervised iterative learning process to understand cell identity within the tissue architecture and to capture single-cell and spatially informed gene characteristics across various datasets. We implemented sparse matrix techniques and advances in distributed graphical processing unit (GPU) training to allow efficient training of FuseMap on the large-scale atlases.

## Benchmarking FuseMap for integrating spatial transcriptomics atlases across technologies

We systematically evaluated FuseMap’s performance using four representative sections with single-cell resolution from similar anterior-posterior locations in image-based Atlases 1 to 4^1-4^, respectively (Methods). The four sections exhibited heterogeneous gene panels, together encompassing 1,998 genes but which consist of only 52 overlapping genes (Extended Data Fig. 1a). First, we evaluated the performance of FuseMap on single-cell embeddings. Unlike scRNA-seq integration methods that only utilized the limited 52 overlapping gene^12-16^, FuseMap effectively incorporates all 1,998 genes across the brain sections. This approach proved beneficial as we observed better corrected batch effects and preserved cellular heterogeneity in the integrated single-cell embedding by FuseMap (Extended Data Fig. 1b-f). We also compared with multi-omic integration methods that can utilize distinct gene panels from each section^16-20^, where FuseMap remained outperformed (Extended Data Fig. 1g-k). To quantitatively benchmark its performance, we generated unified cell-type labels of the four brain sections by transferring the labels of an external annotated scRNA-seq mouse brain atlas (scRNA-seq Atlas 8) onto each section via standard single-cell integration^12^. Using the unified cell-type labels, we assessed single-cell integration performance by biology conservation metrics and batch correction metrics^32^. This test showed that FuseMap achieved the highest overall scores compared to all other methods (Extended Data Fig. 1l,m, Fig. 2a,i-iii).

**Figure 2.**
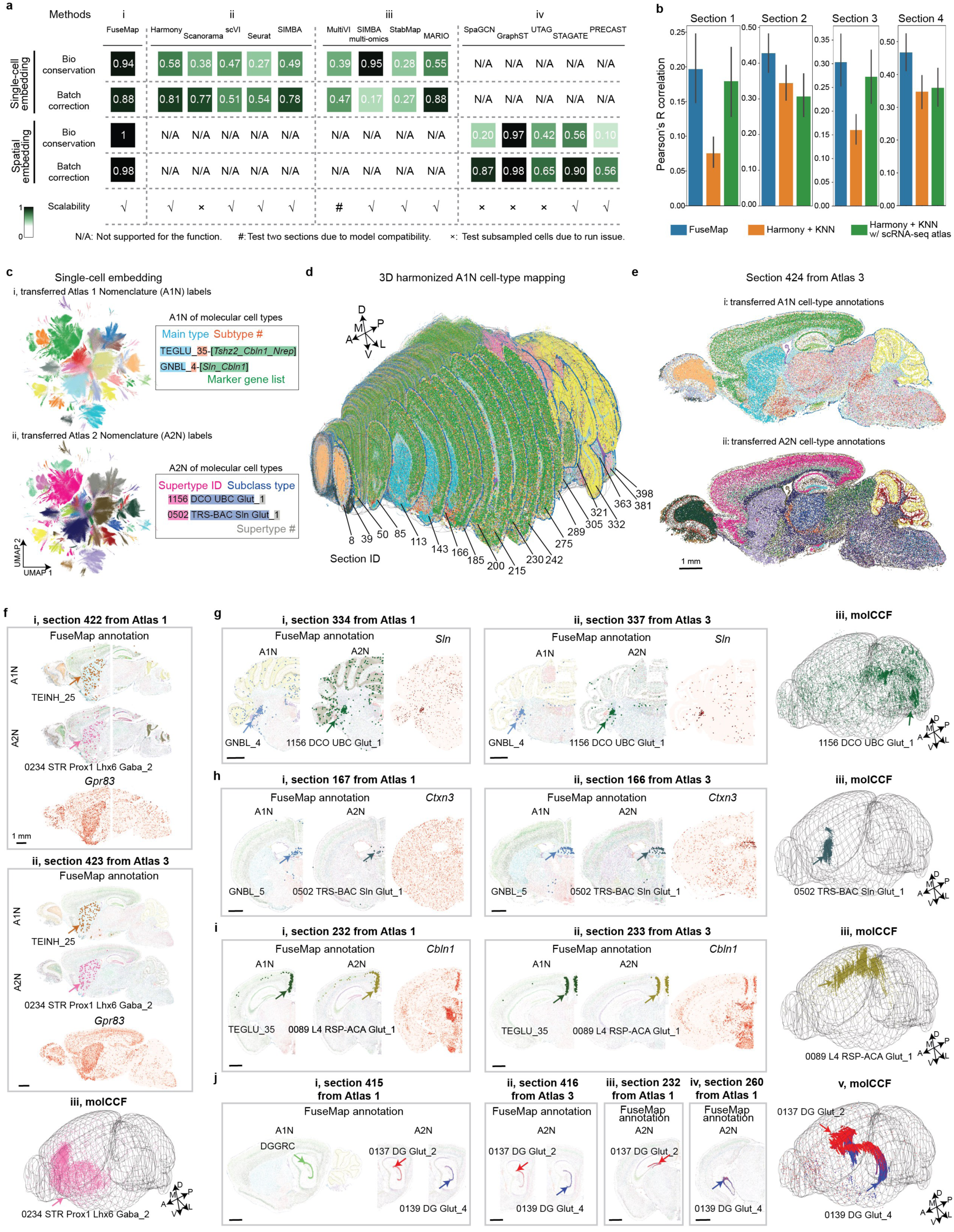
FuseMap-enabled integration and cell-type harmonization across brain atlases. **a,b**, Benchmarking cell-type and tissue-region integration performance (**a**) and gene imputation performance (**b**) of FuseMap with other methods on four representative sections across platforms. **c**, Uniform Manifold Approximation and Projection^43^ (UMAP) of cell-type embedding of randomly sampled one million cells in the atlas corpus, colored by (i) transferred annotation from Atlas 1 nomenclature (A1N), and (ii) transferred annotation from Atlas 2 nomenclature (A2N). **d**, 3D harmonized cell-type mapping of 434 sections (distal hemisphere), with example coronal tissue sections highlighted (proximal hemisphere) in the molCCF. **e**, Representative sagittal section showing (i) transferred cell annotation from A1N, and (ii) transferred cell annotation from A2N. **f-j**, Representative cell subtype correspondence between A1N TEINH_25 and A2N 0234 STR Prox1 Lhx6 Gaba_2 (**f**), A1N GNBL_4 and A2N 1156 DCO UBC Glut_1 (**g**), A1N GNBL_5 and A2N 0502 TRS-BAC Sln Glut_1 (**h**), A1N TEGLU_35 and A2N 0089 L4 RSP-ACA Glut_1 (**i**), and A1N DGGRC and A2N 0137 DG Glut_2 and 0139 DG Glut_4 (**j**), illustrated with distribution in representative tissue sections and the molCCF along with marker gene patterns. Scale bar, 1 mm.

Second, we investigated the performance of FuseMap on spatial embedding. Because FuseMap allows a unified modeling of both single-cell and spatial embedding, it simultaneously enables integration of molecularly defined tissue regions across different sections. Similar to single-cell embedding benchmarking, we generated unified tissue region labels to benchmark FuseMap against existing integration methods^21-25,32^ (Methods). While spatial data integration by previous non-spatial integration methods are largely inaccurate, compared to other spatial integration methods, FuseMap still excelled in removing batch effects and preserving tissue heterogeneity (Extended Data Fig. 2, Fig. 2a,iv).

Third, we examined the capability of FuseMap for imputing unmeasured gene expression in individual sections facilitated by the learned universal gene embedding across sections. We compared FuseMap to two alternative baseline methods^12,33^. One baseline method averages gene expression of *k* nearest neighboring (*k*NN) cells from other sections after integrating single-cell data based on 52 overlapping genes. However, given the limited number of overlapping genes across various image-based spatial transcriptomics sections, this approach would become inaccurate. To compensate this, the other baseline method leverages the scRNA-seq Atlas 8 with extensive transcriptome coverage of 27,998 genes^34^. We evaluated the imputation performance by leveraging a leave-one-out strategy for each intersectional gene and calculating Pearson’s *R* between the imputed and measured expression of the gene (Methods). We found that FuseMap consistently achieved higher correlations than other methods across the four sections (Fig. 2b). To further assess the effectiveness of incorporating spatial information in addition to single-cell information on the imputation performance, we conducted an ablation study by removing the spatial embedding in FuseMap. We observed a corresponding decrease in imputation performance across sections, highlighting the benefits of leveraging both single-cell and spatial embeddings across heterogeneous spatial atlases (Extended Data Fig. 2i).

## FuseMap-enabled universal cell embedding and cell-type annotation harmonization

We then trained a brain FuseMap model on seven spatial atlases of the mouse brain from different technologies^1-7^ encompassing 13.8 million cells from image-based methods and 5.8 million spots from chip-capture-based methods (Methods). These atlases had varying gene panels, spatial resolutions, cell-type compositions, tissue coverages, and transcriptome sparsity, presenting a real-word scenario challenge for unifying organ-wide cell embedding and cell-type annotations. As expected, we observed the learned single-cell embedding effectively eliminated technical variations across different sections and atlases (Extended Data Fig. 3a). We thus tested whether it could harmonize expert-curated cell-type labels across the 13.8 million single cells. By transferring original cell-type annotations from Atlas 1 Nomenclature (A1N) or Atlas 2 Nomenclature (A2N), cells across the entire atlas were accurately categorized (Fig. 2c, Extended Data Fig. 3b,c, Methods). We thus created a harmonized and comprehensive spatial cell-type map of the whole mouse brain, with correspondence between the transferred A1N and A2N (Fig. 2d,e, Extended Data Fig. 3e,d).

Next, we confirmed whether, without any further training, the universal single-cell embedding and cell-type harmonization could enable us to cross-verify region-specific subtypes annotated in original individual atlases^1^. For instance, interneuron subtype A1N TEINH_25^35^ showed alignment with A2N 0234 with consistency across atlases, as supported by their spatial specificity in the striatum and *Gpr83* marker gene expression patterns (Fig. 2f). Similarly, glutamatergic neuroblast A1N GNBL_4 and A1N GNBL_5 correspond with A2N 1156 and 0502, respectively, marked by enriched *Sln* or *Ctxn3* expression. A2N 1156 and A2N 0502 displayed consistent region-specific spatial patterns in the deep cerebellar nuclei or the triangular nucleus of the septum (TRS) within the 3D mouse brain (Fig. 2g,h). Another example was the high correspondence between excitatory neuron subtype A1N TEINH_30 and A2N 0089, specifically localized in the retrosplenial area (RSP) layer 5 (Fig. 2i). These findings confirmed the spatial specificity of these cell types across multiple atlases, which underlies their complex roles in circuitry function and behavior.

Furthermore, cell-type harmonization led to the identification of new cell subtypes previously unannotated in the original publications of individual atlases. For example, with the extensive cell numbers in A2N, we further distinguished subtypes in the A1N dentate gyrus granule cells (DGGRC) and revealed a dorsal-ventral brain gradient from A2N 0137 to A2N 0139 across other atlases and the 3D brain volume (Fig. 2j). In summary, FuseMap harmonizes atlas-level cell types, which not only resolves complex subtype alignments but also enhances cell-type annotations across atlases, potentiating the elucidation of the complex architecture of the brain by the integrated spatial atlas.

## FuseMap-enabled universal spatial embedding toward the molCCF

We next evaluated whether the pretrained brain FuseMap model provided meaningful spatial embedding across various atlases. Despite using the data generated from two different categories of spatial transcriptomics technologies (image-based and chip-capture-based atlases), over 18.6 million cells or spots with prominent technical batch effects, the learned spatial embedding can effectively capture spatially informed gene expression profiles. Based on this, we registered 434 coronal and sagittal spatial transcriptomics tissue sections from ≥11 animals within the Allen Mouse Brain Common Coordinate Framework Version 3 (anaCCFv3)^36-38^, and achieved a comprehensive molecularly defined CCF (molCCF) that offers unified annotations of molecular cells and tissues with their corresponding anatomic location in the anaCCFv3 as detailed below (Supplementary Fig. 1a, Methods).

In our previous publication^1^, we demonstrated the molecular tissue regions defined by spatially resolved gene expression information. As these labels were never used for model training, we used them to validate the quality of the learned embedding. Here, based on the new spatial embedding, we transferred the A1N molecular tissue region labels to all cells or spots across different atlases. This further enabled harmonization of 17 A1N main level molecular tissue regions across seven atlases with 94.0% accuracy across all major brain regions, including the olfactory bulb (OB), cerebral cortex (CTX), cerebral nuclei (CNU), thalamus (TH), hypothalamus (HY), midbrain (MB), hindbrain (HB), cerebellum (CB), fiber tracts (FbTrt), and ventricular systems (VS) (Extended Data Fig. 4a-c, Methods). This demonstrates that the pretrained brain FuseMap model, without any explicit training or labels, identifies comprehensive spatial niche heterogeneities that are shared across 434 tissue samples. Notably, while we achieved an overall high level of correspondence, subtle differences in the transferred annotations in Atlas 1 sections can be observed to update the molecular tissue region segmentation. These updated results demonstrate the advantage of FuseMap in capturing detailed tissue structures in the learned spatial embedding, which were challenging to characterize in the previous *k* or radius nearest neighbor-based approach^1^. For example, in Section 39 from Atlas 1, the main level molecular tissue region annotation of the frontal pole of layer 1 was previously jointly labeled as OB_2 with other OB regions, but is now separated and accurately annotated as L1_HPFmo_Mngs. The refinement is consistent with distribution patterns observed in neighboring sections from other atlases, along with enriched expression of *Myoc* or *Hspb8*, which is further supported by anatomically defined regions in the anaCCFv3^36-38^ (Extended Data Fig. 4d). Other examples included refinements from the original A1N LSX_HY_MB_HB to transferred Hbl_VS in Section 404 and FbTrt in Section 185, all supported by marker genes, consistent spatial distribution in other atlases, and alignment with the anaCCFv3 (Extended Data Fig. 4e,f).

We then explored the sublevel brain molecular tissue regions by hierarchically transferring sublevel A1N molecular tissue region annotations. Since FuseMap expanded the gene-plex from 1,022 to 26,665 with increased cell or spot sampling from 1 to 18 million, we asked whether this integrated atlas could allow us to identify new molecular tissue regions. To answer this, we applied Leiden clustering to each sublevel brain region in this universal embedding and identified molecular tissue regions. We verified the identified molecular tissue regions guided by the following criteria: (i) distinct spatial separation along with region-specific marker genes, (ii) consistent spatial distribution across the entire brain, cross-validated through multiple spatial transcriptomics technologies, and preferably (iii) precise alignment with anatomical definitions outlined by the anaCCFv3^36-38^ (Methods). As a result, we annotated a total of 146 sublevel tissue regions, which were named following original A1N sublevel annotations and the anatomical nomenclature in the Allen Institute Adult Mouse Atlas^38-40^ (Fig. 3a, Supplementary Table 2).

**Figure 3.**
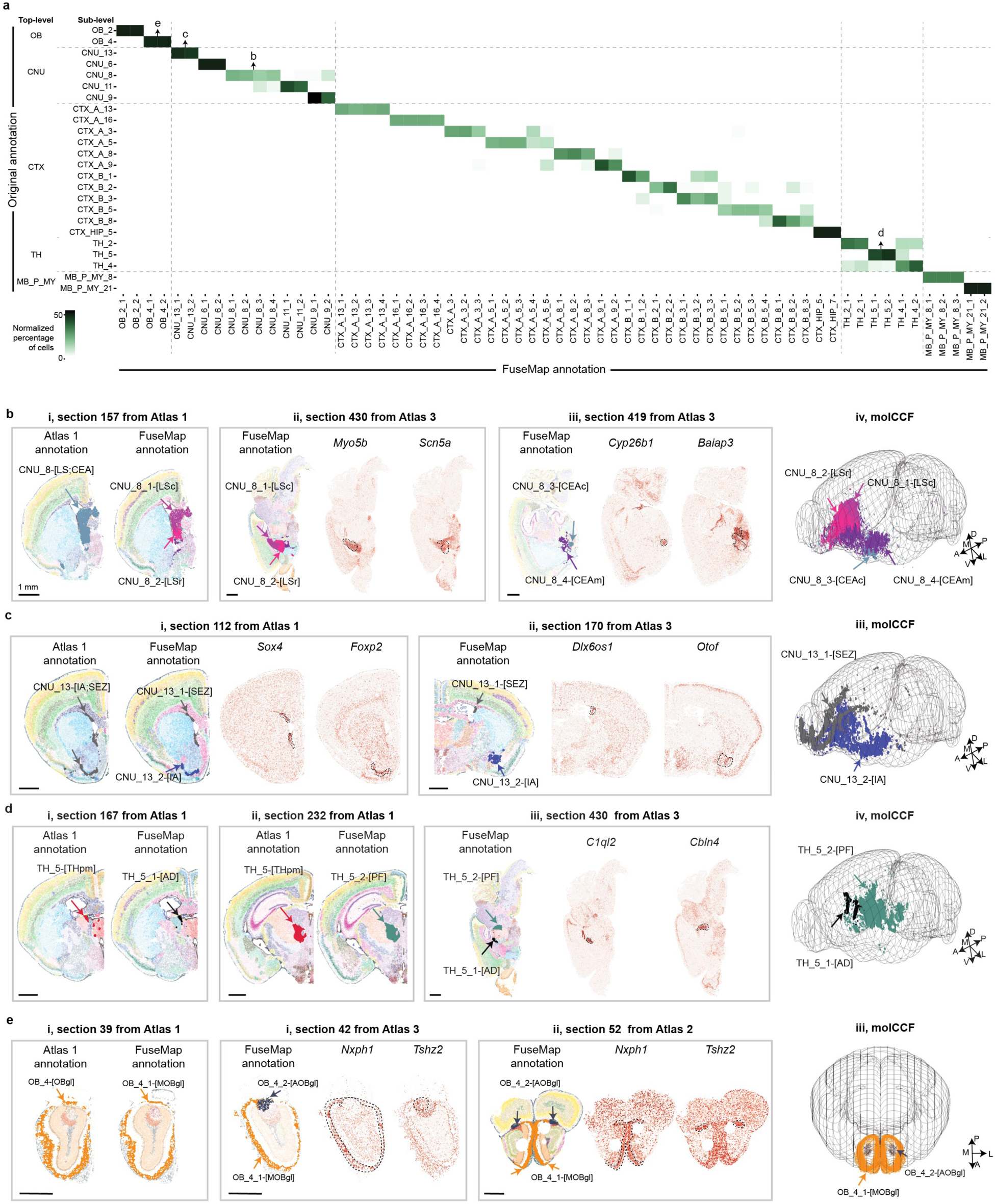
FuseMap-enabled identification of new sublevel molecular tissue regions across the atlas corpus. **a**, Percentage of original A1N sublevel molecular tissue regions (rows) with refined FuseMap annotations (columns). **b-e**, Representative newly identified sublevel tissue regions in A1N CNU_8 (**b**), A1N CNU_13 (**c**), A1N TH_5 (**d**), and A1N OB_4 (**e**), illustrated with distribution in representative tissue sections and the molCCF along with marker gene patterns. LS, lateral septal nucleus; LSc, LS caudal (caudodorsal) part; LSr, LS rostral (rostroventral) part; CEA, central amygdalar nucleus; CEAc, CEA capsular part; CEAm, CEA medial part; IA, intercalated amygdalar nucleus; SEZ, subependymal zone; THpm, thalamus, posterior medial boundary; AD, anterodorsal nucleus; PF, parafascicular nucleus; MOBgl, main olfactory bulb, glomerular layer; AOBgl, accessory olfactory bulb, glomerular layer. Scale bar, 1 mm.

Among the 146 FuseMap-annotated tissue regions, 82 correspond to original annotations (Extended Data Fig. 5a, Methods). For instance, in Atlas 1, we previously identified two distinct regions within the dentate gyrus, A1N CTX_HIP_1-[DGd-sg] and CTX_HIP_2-[DGv-sg]^1^. Using the spatial harmonization by FuseMap, we reproduced the finding by corroborating the existence of these two regions across multiple atlases, marked by additional marker genes *Dgkh* and *Tenm2*, respectively. The spatial mapping in the molCCF further offered complete 3D segmentation of the two molecular tissue regions along the dorsal-ventral axis (Extended Data Fig. 5b). Another example was A1N CNU_12-[TRS] identified in one sagittal section of Atlas 1, showing unique distributions in the coronal sections from other atlases, together unveiling a detailed 3D structure in the molCCF (Extended Data Fig. 5c).

While 82 regions are in line with previous results, benefiting from enlarged cell and gene sampling from multiple atlases, we achieved a more comprehensive molecular tissue-region re-annotation of 64 regions from previously annotated 24 regions by adding 40 previously unannotated tissue regions. First, benefiting from gene expansion from 1,022 to 26,665, FuseMap revealed spatial heterogeneity in areas previously lacking significant molecular differentiation. For example, within A1N CNU_8-[LS;CEA], we identified four sublevel molecular tissue regions, corresponding to the caudal and rostral part of lateral septal nucleus (LSc and LSr), and capsular and lateral part of central amygdalar nucleus (CEAc and CEAm) (Fig. 3b,i). These regions exhibit consistent spatial distribution across different atlases and are characterized by marker genes such as *Myo5b*, *Scn5a*, *Cyp26b1*, and *Baiap3*, respectively, from Atlas 3 (Fig. 3b,ii,iii). Their 3D molecular boundaries are also delineated within the molCCF, aligning with the anaCCFv3 (Fig. 3b,iv). Additionally, in the cerebral cortex, the A1N CTX_A_16-[CLA;EP] region has been further subdivided into four distinct areas, corresponding to (i) the anatomically-annotated claustrum (CLA), (ii) the dorsal part of the endopiriform nucleus (EPd), (iii) the ventral part of EP (EPv) and the medial entorhinal areas of cortical layer 5 (ENTm5), and (iv) the lateral entorhinal areas of cortical layer 5 (ENTl5), each with unique marker genes and spatially segregated in 3D molCCF (Extended Data Fig. 6a). Second, increasing cell numbers from 1 million to 18 million revealed more sublevel tissue regions. For instance, within A1N CNU_13-[IA;SEZ], the subependymal zone (SEZ) became distinguishable with differentially expressed genes *Sox4* and *Dlx6os1*. The newly identified CNU_13_1-[SEZ] region in the molCCF shows a clear rostral migratory stream path from the lateral ventricle to the olfactory bulb ^3^ (Fig. 3c). In the thalamus, the anterodorsal nucleus (AD) is differentiated from the A1N TH_5-[THpm], with enriched *C1ql2* expression (Fig. 3d). Third, since each atlas individually mapped a fraction of the whole-brain volume, the integrated atlas corpus characterized new regions from areas that were previously not sampled in Atlas 1. For instance, in Atlas 1, sampling was limited to the glomerular layer of the main olfactory bulb annotated as A1N OB_4-[OBgl] with insufficient sampling of the glomerular layer of the accessory olfactory bulb; integrated with Atlas 2 and 3, A1N OB_4-[OBgl] was separated into OB_4_1-[MOBgl] and OB_4_2-[AOBgl] (Fig. 3e). Furthermore, we explored new regions in areas that were previously underexplored such as the dentate gyrus polymorph layer (DG-po) and the medial septal complex (MSC) in Atlases 2, 3 and 7 (Extended Data Fig. 6b,c). This demonstrates that the pretrained brain FuseMap model can learn unique tissue regions information through the spatial embedding of multiple atlases.

Finally, comparative analyses of the molCCF and anaCCFv3 further uncovered molecular spatial heterogeneity previously limited by undersampling from individual atlas or not delineated by anatomical structures. For instance, we achieved more accurate molecular tissue regions of piriform area molecular layer (PIR1), pyramidal layer (PIR2), polymorph layer (PIR3), postpiriform transition area layer 2 (TR2), anterior olfactory nucleus layers 1 (AON1) and layers 2 (AON2) within A1N CTX_B_1-3 (Extended Data Fig. 7a-c). Overall, the spatial distribution of these refined molecular regions demonstrated good correspondence with the anaCCFv3^36-38^ along the lateral to medial axis (Extended Data Fig. 7d,e). Notably, two molecular regions CTX_B_3_2-[PIR2l] and CTX_B_3_3-[PIR2m], located in the lateral and medial parts respectively, further subdivided the anatomical PIR2. In addition, in our previous study, we found a discrepancy between molecularly defined and anatomically defined RSP tissue regions^1^. With refined and harmonized tissue annotations across the molCCF, we confirmed that RSP molecular regions including layer 2/3, 5 and 6 align with anatomical RSP towards the anterior of the anterior–posterior axis but have less consensus towards the posterior. Specifically, we refined the originally identified three molecular regions in Atlas 1 including - CTX_A_5, CTX_A_10, and CTX_A_13 - into seven distinct regions as follows: CTX_A_5 to RSPv2/3, RSPd2/3, and presubiculum-postsubiculum (PRE-POST) layers 1/2; CTX_A_10 to RSP5; and CTX_A_13 to RSP6, subiculum ventral part pyramidal layer (SUB-sp), and PRE-POST layer 3 (Extended Data Fig. 8a,b). The refined regions revealed spatial distributions of more detailed molecular RSP tissue regions, with specific markers such as *Gpr101* and *Adgrd1* identifying SUB-sp and PRE-POST3, respectively (Extended Data Fig. 8c,d). Furthermore, we identified more comprehensive distributions in previously unexamined areas from individual atlas along the anterior to posterior axis. This finding also reveals that, in the interconnected region of RSP and PRE/POST, transcriptional signatures of different laminar layers display significant heterogeneity, which can be distinguished at the main level of clustering. However, tissue regions in this area that define functions for RSP, SUB, PRE, and POST are distinguished at a sublevel tissue region analysis, highlighting the complexity and understanding of molecular and anatomical correlations.

In conclusion, FuseMap achieved universal spatial embedding and atlas-level harmonization of molecular brain regions at both main level and sublevel (Supplementary Fig. 1b, Methods).

## FuseMap-enabled transcriptome-wide spatial gene expression imputation at single-cell resolution

Considering the varying gene panels and spatial resolution across multiple spatial atlases including targeted gene panels in Atlases 1 to 5 with single-cell resolution and non-targeted transcriptomics Atlases 6 and 7 (Extended Data Fig. 9a), we sought to impute transcriptome-wide spatial gene expression combining all cells and spots to restore spatially resolved transcriptome at single-cell resolution. The pretrained brain FuseMap model embedded all 26,665 genes into a low-dimensional space that encoded the characteristics of their spatial expression patterns. We observed that the learned gene embeddings were not only robust to technological platforms but also encompassed both shared and distinct genes without atlas-specific bias (Fig. 1d,iii). We thus performed coupled matrix multiplication of the universal gene and single-cell embedding to impute single-cell or single-spot expression of all genes (Extended Data Fig. 9b, Methods). To assess the imputation accuracy on a tissue section, we leveraged adjacent tissue sections in the molCCF, which share most similar patterns of spatial cell atlas and are typically from a different atlas with varying gene panels. Through registration to the molCCF and binning cells or spots with a unified square size, we could estimate correlations between the two sections in three groups of comparisons (Extended Data Fig. 9c,d, Methods): (i) Group 1, between measured expressions of shared genes as ground truth, (ii) Group 2, imputed versus measured expressions of shared genes, and (iii) Group 3, imputed versus measured expressions of distinct genes.

In comparing adjacent tissue sections across two different image-based spatial transcriptomics atlases, e.g. Atlas 1 and Atlas 3, we observed higher average correlations for shared genes’ imputed expression (Group 2) compared to the measured expression (Group 1), suggesting reduced technical variance in imputed expression. Furthermore, the correlations for shared genes and distinct genes were comparable (Groups 2 versus 3), exceeding the benchmark set by Group 1, indicating FuseMap’s effectiveness in predicting expressions of unmeasured genes (Fig. 4a). As an example, the distribution of correlations across genes between representative Section 208 from Atlas 1 to Section 211 from Atlas 3 supports the result (Fig. 4b).

**Figure 4.**
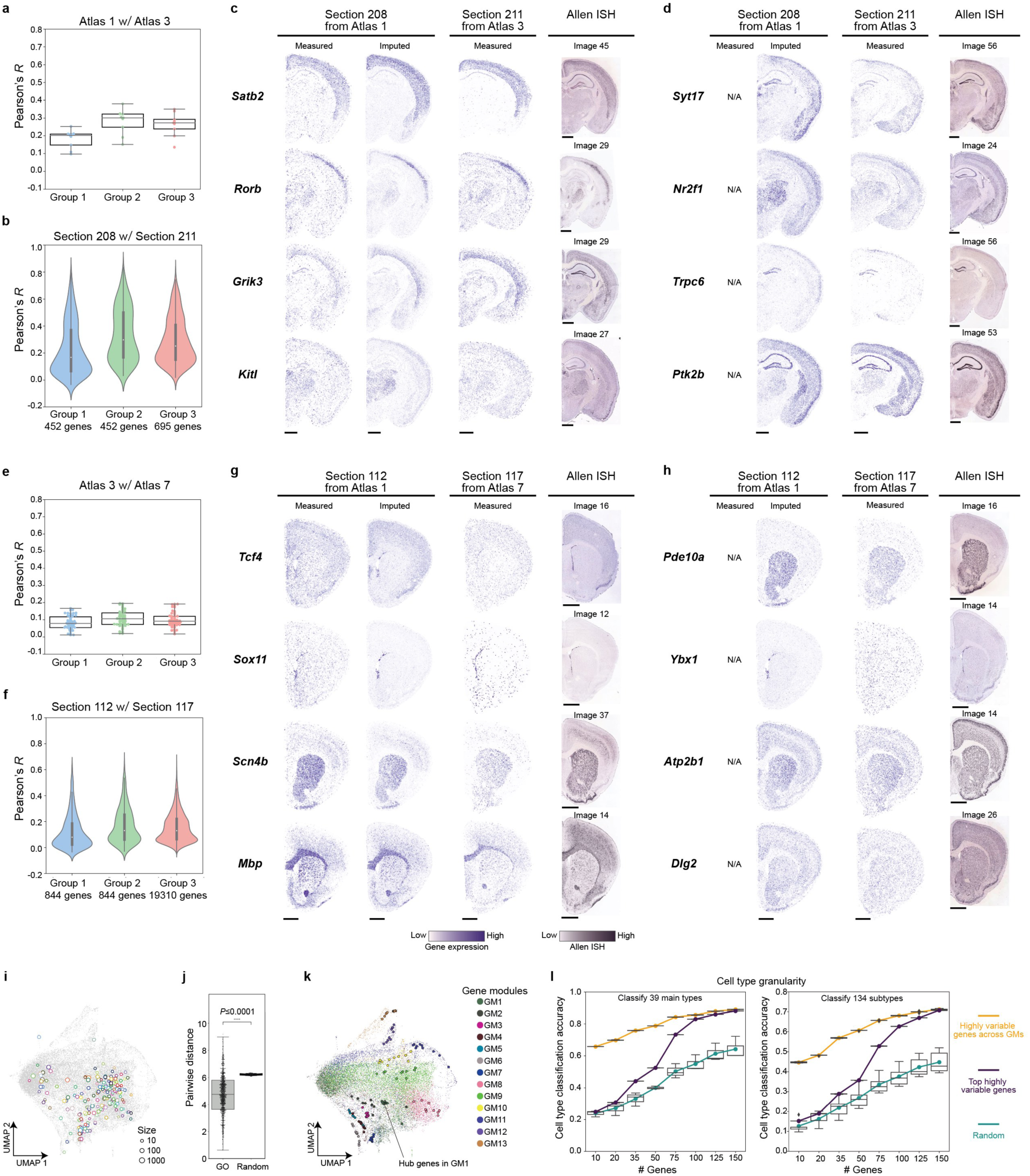
FuseMap-enabled spatial gene imputation and gene list selection. **a,** Average Pearson’s *R* of 10 neighboring sections between Atlas 1 and Atlas 3. From left to right: measured expressions of shared genes (Group 1), imputed versus measured expressions for shared genes (Group 2), and imputed versus measured expressions for distinct genes (Group 3). **b**, Violin plots of Pearson’s *R* between Section 208 and 211. **c**,**d**, Representative measured and imputed spatial gene expression maps on Section 208 from Atlas 1, Section 211 from Atlas 3, and Allen Mouse Brain ISH expression. Each dot represents a cell colored by the expression level of a gene. Shared (**c**) and distinct (**d**) genes are included. **e,** Same as (**a**) but between Atlas 3 and Atlas 7 with 58 neighboring sections. **f**, Same as (**b**) but on Section 112 and 117. **g**,**h**, Same as (**c**,**d**) but on Section 112 from Atlas 1, Section 117 from Atlas 7, and Allen Mouse Brain ISH expression. Shared (**g**) and distinct (**h**) genes are included. **i**, Centers of genes within the same protein family on UMAP of the gene embedding. 199 protein families with significance were shown. The size reflects the number of genes. **j**, Boxplot of UMAP distance between genes within the same protein families and random selected genes. Statistical test *P* < 0.0001, Mann-Whitney-Wilcoxon test. **k**, UMAP of gene embedding colored by gene modules (GM). Hub genes within each GM are highlighted. **l**, Cell type classification accuracy for gene panels selected by different strategies within the scRNA-seq Atlas 8. Accuracy is evaluated on 39 (left) and 134 (right) cell type granularity. Scale bar, 1 mm.

The spatial patterns at single-cell resolution displayed consistent patterns between imputed and experimental results of shared genes, including *Satb2* as an excitatory neurons marker, *Rorb* as a layer 4 cortical neurons marker, *Grik3* as functional marker involved in excitatory neurotransmission, and *Kitl* as an adhesion molecule (Fig. 4c). In addition, we observed accurate imputations for originally unmeasured genes in Section 208, such as *Syt17* for synaptic vesicle release, *Nr2f1* involved in neuronal development, *Trpc6* as a calcium channel mainly expressed in dentate granule cell, and *Ptk2b* as a focal adhesion kinase involved in cell adhesion and signaling (Fig. 4d). Meanwhile, we benchmarked with the Allen In Situ Hybridization (ISH) database^41^ for individual genes, confirming the accuracy of FuseMap in predicting gene expression across molecular resolved tissue atlas.

We extended our analysis to encompass the entire mRNA repertoire through chip-capture-based spatial transcriptomics atlases. We observed consistently higher averaged correlations in Group 2 over Group 1, and comparable performance between Group 2 and 3 across adjacent tissue sections between image-based Atlas 3 and chip-capture-based Atlas 7 (Fig. 4e). Furthermore, the analysis of Section 112 from Atlas 1 and Section 117 from Atlas 7 revealed a correlated distribution of 844 shared and 19,310 distinct genes, illustrating our findings (Fig. 4f). Notably, the imputed expression in Section 112 from Atlas 1 captured the spatial expression of both shared genes, e.g., *Tcf4*, *Sox11*, *Scn4b*, and *Mbp*, and distinct genes, e.g., *Pde10a*, *Ybx1*, *Atp2b1*, and *Dlg2,* at single-cell resolution, which aligned closely with corresponding ISH images^41^ (Figs. 4g,h). However, overall we noticed lower correlations between image- and chip-capture-based atlases compared to those between image-based atlases. This may be explained by limitations in near-single-cell resolution and gene detection sensitivity in Atlas 7 as observed in Section 117.

## Spatial and single-cell informed selection of a targeted gene panel

We next investigated the gene-to-gene relationships within the learned gene embedding. Utilizing a PANTHER dataset with classification of complete protein-coding gene families in mouse genomes^42^, we examined the distribution of genes grouped within the same protein family or gene ontology category in the gene embedding visualized via Uniform Manifold Approximation and Projection (UMAP)^43^ (Supplementary Fig. 2, Methods). We found pairwise distances between genes from the same protein families significantly lower than distances of randomly sampled gene pairs on the UMAP (Fig. 4i,j, statistical test *P* < 0.001, Mann-Whitney-Wilcoxon test). This suggests that the learned gene embedding provides a comprehensive view of modular characteristics of genes corresponding to functional biological pathways.

Therefore, we further utilize the learned gene embedding to identify an effective gene panel (i.e., a small number of highly informative target genes), which is critical for the design of most image-based spatial transcriptomics experiments^44^. Specifically, we compared gene embedding-guided with non-guided approaches to identify panels of 10 to 150 marker genes from the scRNA-seq Atlas 8^34^. For non-guided approaches, we used a randomly chosen gene panel or a standard panel of top highly variable genes^45^. To leverage the learned gene embedding, we grouped all genes into 13 modules. Then, from each module, we selected a panel of highly variable genes, ensuring an even distribution across all gene modules, even when these genes did not rank as the most variable (Fig. 4k, Methods). As a benchmark, we employed selected gene panels to classify originally defined transcriptomic cell types. The classification accuracy indicates the discriminative power of the gene panels in distinguishing cell types, thereby simulating the performance in a corresponding spatial transcriptomics experiment. The comparisons revealed that the gene embedding-guided selection consistently showed higher classification accuracy over non-guided approaches in both 39-cell-type and granular 134-cell-type taxonomies (Fig. 4l). These results demonstrate the effectiveness of using the universal gene embedding in gene panel selection for targeted spatial transcriptomics experiments.

## Spatially informed cell identity and cell-cell interactions

With FuseMap directly linking transcriptional profiles with spatial information, we wondered whether the learned single-cell embedding could discern subtly different cell subtypes influenced by distinct microenvironments. We observed a clear spatial segregation across distinct brain regions in single-cell embedding of three major A2N mature oligodendrocyte (MOL) types (Supplementary Fig. 3a,i,ii). Further clustering divided each A2N MOL type into two spatially distinct groups: one associated with white matter (WM) and the other with grey matter (GM), consistent with a recent study^46^ (Supplementary Fig. 3a,iii, Methods). Analysis of imputed gene expression revealed upregulation of *Lpar1* and *Zeb2* in the WM groups, suggesting functional roles in neuronal myelination (Supplementary Fig. 3b,c), while *Gad1* and *Syt4* were upregulated in the GM groups, potentially associated with synaptic transmission and signaling roles^46^ (Supplementary Fig. 3d,e). The spatial distribution of these WM and GM groups exhibited distinct patterns with differential gene expression across Atlas 1, 2, and 3 (Supplementary Fig. 3f-h).

Moreover, within the molCCF, the unified molecular cell types, molecular brain regions and transcriptome-wide expression enables us to explore cell-cell interactions arising from soma contact, paracrine signaling or other short-range interactions across different brain atlases. Following previous analysis of comparing observed proximity frequencies to a permutation model of random chance^3^, we identified proximal A1N cell-type pairs in specific FuseMap-annotated molecular brain regions (Methods). First, we examined proximal cell-type pairs within each main level molecular brain region and found that the cell–cell adjacency patterns were overall consistent across Atlas 1, 2 and 3 from different technological platforms (Extended Data Fig. 10a). Meanwhile, we observed a higher number of total proximal cell-type pairs in Atlas 3 (2,374) compared to 1,623 in Atlas 1 and 1,529 in Atlas 2, potentially due to the larger tissue coverage of approximately 6.9 million cells in the molCCF from Atlas 1 compared to 1.1 million from Atlas 1 and 2.0 million from Atlas 2. Therefore, we analyzed altogether over 10 million cells in the molCCF across atlases and identified a total of 2,951 proximal cell-type pairs. Additionally, we analyzed sublevel molecular region-specific cell-cell interactions. For example, we found significant interactions between astrocytes and non-glutamatergic neuroblasts in the subependymal zone CNU_13_1-[SEZ], consistently across Atlas 1, 2 and 3 (Extended Data Fig. 10b-h). Since neuroblasts migrating from the SEZ to the olfactory bulb interact with astrocytic cells along the rostral migratory stream, the observed interactions in the olfactory bulb might be related to the interactions between neuroblasts and astrocytes in the rostral migratory stream^3,47,48^.

## New query data by transfer learning

By transfer learning, the pretrained FuseMap model on reference data can map and annotate new query data with cell-type and tissue-region labels based on the cell embeddings (Fig. 1e, Methods). First, we benchmarked the performance of reference mapping and annotation transfer using previous benchmark sections from different atlases with unified cell-type labels (from ‘Benchmarking FuseMap for integrating spatial transcriptomics atlases across technologies’ section). In a basic setting, one section served as the reference to train a FuseMap model, while another section as query data to be annotated. Comparing predicted labels with unified labels, we observed high accuracy across standard metrics compared to existing annotation transfer methods^49-56^ (Supplementary Fig. 4a). Additionally, we tested a larger reference setting, with three sections as reference and one as query. Interestingly, we observed a scaling effect in FuseMap and one transformer-based method, TOSICA^52^, where pretraining on the larger reference improved performance on annotation transfer to query data (Supplementary Fig. 4a). In contrast, the performance of other annotation methods decreased in the larger reference setting. This finding highlights the efficacy of FuseMap in generalizing and learning reference knowledge across atlases. Next, we evaluated FuseMap on atlas-level annotation transfer using Atlas 1 and Atlas 2 as reference data. We transferred cell-type labels to four leave-out Atlas 2 sections spanning from anterior to posterior across the brain, achieving predicted cell types that matched original annotations with 94.7% accuracy (Supplementary Fig. 4b). Furthermore, we transferred cell-type labels to query data collected from a different animal, unseen in the reference, and from a sagittal view, achieving accurate annotations with 89.1% accuracy (Supplementary Fig. 4c). Meanwhile, we transferred molecular tissue region labels to query data. Transferred region labels aligned well with original annotations, except one region, ENTm, which was not included in reference Atlas 1 coronal sections and further identified in the spatial embedding (Supplementary Fig. 4d). After refinement, the tissue-region annotation accuracy reached as high as 89.9% (Supplementary Fig. 4e).

We then applied the pretrained brain FuseMap model to annotate other brain datasets outside the training Atlases 1 to 7 (Methods). We first applied to a STARmap PLUS query dataset, which encompasses a mouse coronal hemi-brain slice with 5,413 genes^57^. With transfer learning, FuseMap mitigated batch effects between reference and query in the single-cell embedding (Supplementary Fig. 5a). In transferring the cell-type annotations, we observed consistent spatial distributions between main level original and transferred cell types (Fig. 5a, Supplementary Fig. 5b). Furthermore, sublevel transferred annotations extended original 38 subtypes to 143 detailed subtypes (Supplementary Fig. 5c). Specifically, we analyzed telencephalon projecting excitatory neurons where the transferred subtypes were more accurate than the original annotations, with layer-specific distributions (Fig. 5b, Supplementary Fig. 6). In addition, FuseMap also generated an integrated spatial embedding and transferred a detailed taxonomy of molecular brain region annotations at the main- and sublevels (Fig. 5c, Supplementary Fig. 5d,f). The transferred main level molecular brain regions correspond closely with the original anatomically defined regions (Supplementary Fig. 5e), while the transferred sublevel molecular brain regions further delineate finer brain structures (Fig. 5e). For example, sublevel regions in isocortex and hippocampal regions resembled the laminar organization of cerebral cortical layers and CA1-3, and DG (Fig. 5d). We then investigated multi-batch mapping by applying to two MERFISH sections of the mouse primary motor cortex targeting 256 genes^58^. Similarly, we observed consistent integration between reference and query datasets and the transferred annotations showed clearly layered spatial distributions for cell types and tissue regions, confirming the advantage of our transfer approach to conventional manual labeling (Fig. 5f-i, Supplementary Fig. 5g-i).

**Figure 5.**
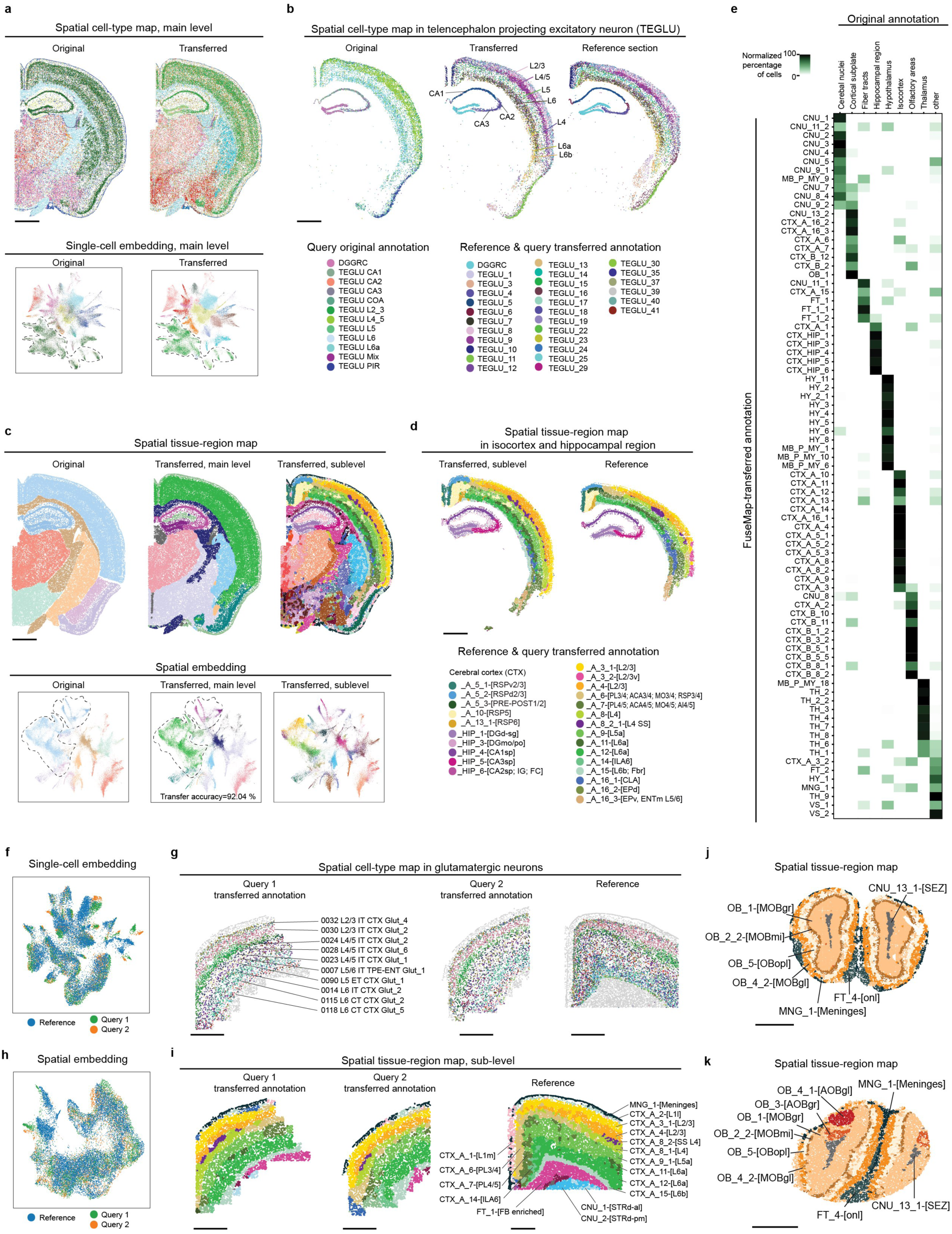
FuseMap-enabled cell type and tissue region annotation transfer to new query data. **a**, Spatial cell-type maps (upper row) and UMAP of cell-type embedding (lower row), colored by main level original (left) and transferred (right) annotations on STARmap PLUS query sample. **b**, Spatial cell-type maps in TEGLU, colored by sublevel original (left) and transferred (middle) annotations on STARmap PLUS query sample and FuseMap (right) annotations on a reference sample. **c**, Spatial tissue-region maps (upper row) and UMAP of tissue-region embedding (lower row), colored by original (left), main level transferred (middle), and sublevel transferred (right) annotations on STARmap PLUS query sample. **d**, Spatial tissue-region maps in isocortex and hippocampal region, colored by sublevel transferred (left) annotations on 5,413-gene STARmap PLUS query sample^57^ and FuseMap (right) annotations on a reference sample. **e**, Percentage of original tissue regions (columns) with transferred sublevel tissue annotations (rows). **f**,**h**, UMAP of cell-type (**f**) and tissue-region (**h**) embedding colored by the dataset, including one reference sample and two MERFISH query samples. **g**, Spatial cell-type maps in glutamatergic neurons, colored by sublevel transferred (left, middle) annotations on two 256-gene MERFISH query samples^58^ and FuseMap (right) annotations on a reference sample. **i**, Spatial tissue-region maps, colored by sublevel transferred (left, middle) annotations on two MERFISH query samples and FuseMap (right) annotations on a reference sample. **j**,**k**, Spatial tissue-region maps of transferred sublevel annotation in Stereo-seq (**j**) and Slide-seq V2 (**k**) query datasets^6,59,60^. IA, intercalated amygdalar nucleus; MOBgr, main olfactory bulb, granule layer; OBmi, olfactory bulb, mitral layer; OBopl, main olfactory bulb, outer plexiform layer; MOBgl, main olfactory bulb, glomerular layer; onl, olfactory nerve layer of main olfactory bulb; AOBgl, accessory olfactory bulb, glomerular layer; AOBgr, accessory olfactory bulb, granular layer. Scale bar, 1 mm.

Furthermore, we applied the pretrained brain FuseMap model to chip-capture-based spatial transcriptomics platforms by mapping two sections of the mouse olfactory bulb from Slide-seq V2^59^ and Stereo-seq^6,60^. Despite significant batch effects, FuseMap successfully transferred common and unique tissue structures across the two datasets. (Fig. 5j,k). Six discrete layers of OB shared by both sections are spatially organized from the innermost to the outermost layers of the mouse olfactory bulb, including the SEZ in CNU_13_1, the granule layer in OB_1, the mitral layer in OB_2_2, the outer plexiform layer in OB_5, the glomerular layer in OB_4_2, and the olfactory nerve layer in FT_4. Furthermore, FuseMap annotated two unique spatial structures in the Slide-seq V2 covering the assessory olfactory bulb: the glomerular layer in OB_4_1 and the granular layer in OB_3 (Fig. 5k).

## Discussion

We introduce FuseMap, a deep-learning framework for spatial transcriptomics that models cell identities and tissue structures by incorporating both single-cell expression and spatial contexts. FuseMap learns single-cell and spatial embeddings, paired via unique physical tissue coordinates, and introduces a universal gene embedding consolidating various gene panels across technological platforms. Comparative analyses show that FuseMap excels in integrating single-cell embeddings, spatial embeddings, and gene imputation across spatial transcriptomics atlases, outperforming existing methods (Fig. 2a).

We trained FuseMap on multiple spatial transcriptomics brain atlases encompassing over 18.6 million cells or spots and 26,665 genes with a self-supervised learning approach (Fig. 1). The pretrained brain FuseMap model generates universal embeddings for cells and genes. With the single-cell embedding, we systematically harmonized cell-type annotations across atlases into two hierarchical nomenclatures: A1N, with 26 main and 191 sublevels, and A2N, with 33 main classes, 332 subclasses, 1,177 sub-classes, and 4,133 clusters. This harmonization facilitates cross-validation and refinement of transcriptomic distinct cell subtypes (Fig. 2). With the spatial embedding, we explored molecular brain regions delineated through spatial niche gene expression profiles. Notably, the integration across atlases significantly increased cell and gene sampling, leading to the identification of 40 new molecular tissue architectures, totaling 17 main levels and 146 sublevels of molecular brain structures (Fig. 3). With the gene embedding, we expanded to 26,665 transcriptome-wide spatial gene expressions and enhanced targeted gene panel selection for image-based spatial transcriptomics experiments (Fig. 4). Altogether, we established the molCCF with individual RNA molecules, single cells and tissue regions, providing a roadmap for investigating transcriptome-scale gene expression patterns and cell-type diagrams within brain architecture. Using the molCCF, we identified molecular spatial heterogeneity not delineated by anatomical structures, such as in piriform area pyramidal layer and retrosplenial area (Extended Data Figs. 7,8). We also investigated distinct subtypes of mature oligodendrocytes across white matter and grey matter regions, suggesting their varied functional roles in myelination and synaptic transmission (Supplementary Fig. 3). Moreover, we identified consistent cell-cell interactions across different brain atlases (Extended Data Fig. 10).

Furthermore, the pretrained brain FuseMap model enables explorations into the brain’s heterogeneity and tissue complexity efficiently by accurately mapping and annotating query datasets (Fig. 5). In the future, FuseMap has the potential to link transcriptomic identities with other spatial multi-omics approaches, such as epigenomics, translatomics, proteomics, and functional imaging and recording^57,61,62^, which will enrich our understanding into the distinct functional roles of individual cells in the brain. Finally, FuseMap is also applicable to virtually any types of spatial tissue atlases.

## Methods

### FuseMap framework

#### Data overview

FuseMap is a deep learning framework for integrating single-cell or single-spot and spatially resolved gene expression profiles across heterogenous atlases. Suppose that there are 𝑉 different tissue sections, each with a distinct gene set 𝒯^𝑣^, 𝑣 = 1,2, …, 𝑉. Let 𝑛^𝑣^ denote the number of cells and |𝒯^𝑣^| denote the number of genes in the 𝑣-th tissue section. Let 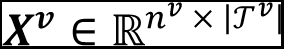 denote the gene expression data where its 𝑖-th row 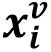 corresponds to the expression of cell or spot 𝑖 in 𝑣-th tissue section. Let 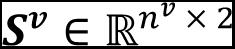 denote the spatial data where its 𝑖-th row 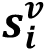 corresponds to the 2D spatial coordinate of cell or spot 𝑖 in 𝑣-th tissue section.

#### *Data* preprocess

For spatial transcriptomics data in 𝑣-th tissue section, its gene expression matrix 𝑋^𝑣^is normalized, log-transformed and scaled using standard processing in Scanpy. To utilize the spatial coordinate 𝑆^𝑣^, we convert it into an undirected neighborhood graph 𝐺^𝑣^ by Delaunay triangulation. In the graph 𝐺^𝑣^, 𝐴^𝑣^ ∈ 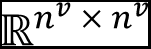 is the adjacency matrix where values are set to 1 if there is connection between adjacent cells, and 0 for no presence of an edge. Next, to account for self-connections in the graph, we add an identity matrix of the same dimension to 𝐺^𝑣^. To normalize 𝐴^𝑣^, we compute the degree of each node, which is the sum of each row in 𝐴^𝑣^, and construct a diagonal matrix 𝐷^𝑣^with diagonal entries being the inverse square root of the degree. The normalized adjacency matrix is obtained by multiplying 𝐴^𝑣^ with 𝐷^𝑣^ from both sides, ensuring symmetric normalization. For simplicity we still represent the normalized adjacency matrix as 𝐴^𝑣^. The compressed sparse row or coordinate format matrix format was chosen to store 𝑋^𝑣^and 𝐴^𝑣^for the efficient memory-saving properties and arithmetic operations, especially suitable for large graphs, as it stores only non-zero elements.

#### Model architecture

FuseMap models the observed transcriptomic expression profiles and spatial contexts of cells generated by a unified low-dimensional latent representation (i.e., cell embedding). Thus, FuseMap approaches a 𝑘 dimensional cell-type embedding 𝑧_𝐶_ ∈ ℝ^𝑘^ from measured transcriptomic modality.

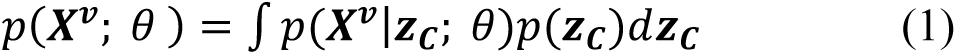

where 𝑝(𝑧_𝐶_) is the prior distribution of the latent representation, 𝑝(𝑋^𝑣^|𝑧_𝐶_; θ) is learnable generative distributions and θ is learnable parameters in the decoders. The cell-type embedding 𝑧_𝐶_ represents the common cell states and is observed from transcriptional modality.

FuseMap then approaches a tissue-region embedding 𝑧_𝑇_ D ℝ^𝑘^ from the cell-type embedding 𝑧_𝑐_ that encodes measured gene expression profiles and the adjacency matrix 𝐴^𝑣^ that encodes measured spatial locations. Specifically, we utilize a graph convolutional network (GCN) as encoder to learn 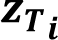 for cell or spot 𝑖 by iteratively aggregating the cell-type embeddings of its neighbors. A 𝑙-th layer representations is:

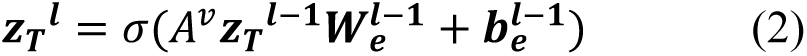

where 𝐴^𝑣^ represents the normalized adjacent matrix. 𝘸_𝑒_, 𝑏_𝑒_, and 𝜎(·) denote a trainable weight matrix, a bias vector, and a nonlinear activation function. 𝑧_𝑇_^𝑙–1^ denotes the (𝑙 − 1)-th layer output representation. 𝑧_𝑇_^0^ is set as the learned cell-type embedding 𝑧_𝐶_, and 𝑧_𝑇_ as the final output of the encoder.

After that, the tissue-region embedding 𝑧_𝑇_ are fed into a decoder to reverse them back into the learned cell-type embedding 𝑧_𝑐_. The decoder adopts a symmetric architecture to reconstruct the cell-type embedding.

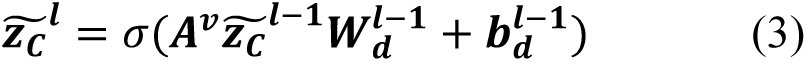

where 𝘸_𝑑_, 𝑏_𝑑_, and 𝜎(·) denote a trainable weight matrix, a bias vector, and a nonlinear activation function. 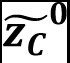 is set as the tissue-region embedding 𝑧_𝑇_ and 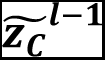 denotes the reconstructed cell-type embedding at the (𝑙 − 1)th layer.

FuseMap initializes a gene embedding matrix as a trainable parameter of the model, denoted as 𝐸 ∈ 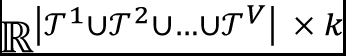 where |𝒯^1^ ∪ 𝒯^2^ ∪ … ∪ 𝒯^𝑉^| is the number of all genes across all tissue sections and 𝑘 is the vector dimension corresponding to the latent space dimensionality of cell embeddings 𝑧_𝐶_ and 𝑧_𝑇_. Its j-th row 𝑒_j_corresponds to the latent embedding of gene j, shared across all atlases. 𝐸 is initialized with the Xavier uniform initialization to populate the gene embedding matrix with values and set the stage for an efficient training process. During training, for a set of 𝒯^𝑣^genes in the 𝑣-th tissue section, we refer their latent gene embedding as 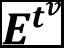.

We model 𝑋^𝑣^ as generated by the combination of latent gene embedding 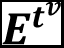 and the latent cell-type embedding 𝑧_𝐶_. The model likelihood (1) can thus be written as:

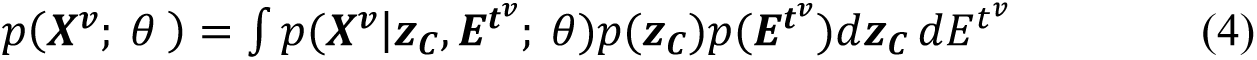

where 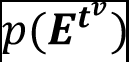 are the distribution of the gene embedding. The gene embeddings 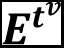 confer semantic meanings for the cell embedding space. Thus, the shared gene embeddings link the independently parameterized and learned cell embeddings for each tissue section and enable them to have consistent semantic meanings.

#### Loss function

The discriminator loss measures how discriminative different tissue sections are in the cell-type and tissue-region embeddings. The loss function ℒ_𝐷𝑖𝑠_ is the sum of the multiclass cross-entropy losses for both the cell-type and tissue-region embeddings.

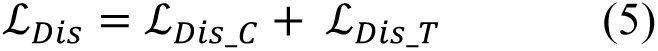

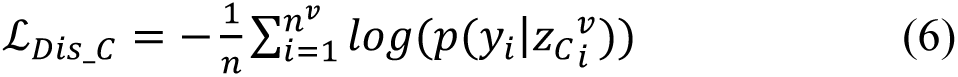

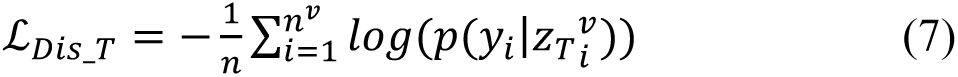

where 𝑦_𝑖_ represents the tissue section source identifier for each cell.

The autoencoder loss is the sum of two components, the reconstruction loss to raw gene expression data and the Kullback–Leibler (KL) divergence.

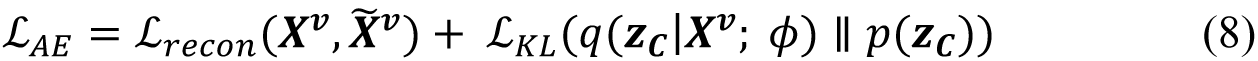

where 𝑋^𝑣^ is the raw gene expression data and 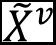 is the reconstructed gene expression data from the autoencoder. 𝑞(𝑧_𝐶_|𝑋^𝑣^; 𝜙)is the variational posterior distribution of the cell-type embedding 𝑧_𝐶_ given the input 𝑋^𝑣^, parameterized by 𝜙. 𝑝(𝑧_𝐶_) is the prior distribution of the cell-type embedding, assumed as a standard normal distribution 𝒩(0,1).

The reconstruction loss ℒ_𝑟𝑒𝑐𝑜𝑛_ is the mean squared error (MSE) between 𝑋^𝑣^ and 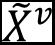, defined as:

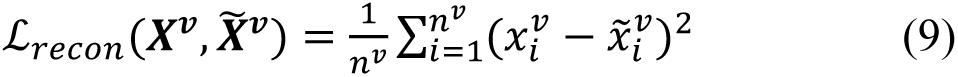

where 𝑛^𝑣^ denote the number of cells in the 𝑣-th tissue section.

The KL divergence ℒ_𝐾𝐿_ is computed between the variational posterior 𝑞(𝑧_𝐶_|𝑋^𝑣^; 𝜙) and the prior 𝑝(𝑧_𝐶_), summed over all dimensions and averaged across the tissue sections. It is normalized by the number of genes |𝒯^𝑣^|. The KL divergence quantifies the difference between the two distributions and acts as a regularization term.

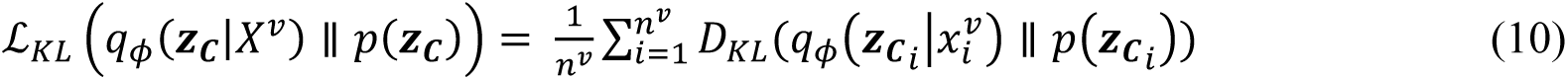

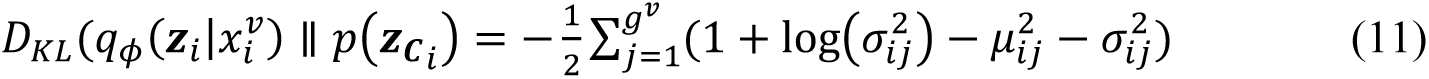

where 𝜇_𝑖j_ and 𝜎_𝑖j_ are the mean and standard deviation of the *j*-th gene in the *i*-th cell’s cell-type embedding, respectively.

The learning process for multiple spatial transcriptomics datasets entails adversarial optimization of the minimizing discriminator loss ℒ_𝐷𝑖𝑠_ to ensure integration of tissue samples, and generator loss trained in the opposite direction to approach correct latent cell embeddings.

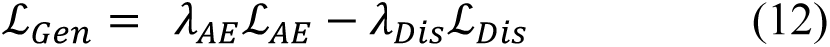

where 𝜆_𝐷𝑖𝑠_ and 𝜆_𝐴𝐸_ are weighting factors.

#### Training procedure

The learning process is a self-supervised iterative learning process, eliminating the need for labeled data. Each spatial transcriptomics sample’s unique characteristics are modeled using dedicated autoencoders and graph autoencoders, forming a sample-specific probabilistic generative model. During the model training process, 15% of the cells are hold out as the validation set. Training is designed to early stop if there was no improvement in the validation loss for consecutive epochs. The training procedure includes a pretraining phase, a weighted adversarial alignment as employed in previous research, and a final training phase that utilized cell-specific weights to ensure effective integration for cases when cell type compositions vary intrinsically.

#### Transfer learning

For a pretrained FuseMap model on reference datasets, we can map and annotate a query dataset through transfer learning. Instead of training the model from scratch, the pretrained model is fine-tuned on the query dataset. For cell embedding, FuseMap freezes the parameters in pretrained autoencoders and graph autoencoders for reference samples and only trains on dedicated autoencoders and graph autoencoders of query dataset. For gene embedding, FuseMap freezes the learned gene embedding for genes in the reference dataset and only updates new distinct genes in the query dataset.

### Benchmark integration performance

To evaluate integration performance, we used four representative sections with single-cell resolution from Atlases 1 to 4 respectively^1-4^. Section 1 from Atlas 1 ‘Well 11’ includes 43,341 cells, 1,022 genes. Section 4 from Atlas 2 ‘C57BL6J-638850.38’ includes 119,927 cells, 500 genes. Section 3 from Atlas 3 ‘col_slice37’ includes 44,959 cells, 1122 genes. Section 2 from Atlas 4 ‘S2R1’ includes 83,546 cells, 483 genes.

To evaluate the performance of integration, we manually labelled harmonized cell types and tissue regions as ground truth. For harmonized cell types on four sections, we integrated each section individually with the scRNA-seq Atlas 8 by Harmony^12^ and transfer scRNA-seq cell types to spatial transcriptomics data using *k*-nearest neighbors (KNN, here *k*=50). For harmonized tissue regions on four sections, we spatially clustered each section by STAGATE^24^ and manually aligned the identified coherent tissue regions labels. Therefore, the four representative sections were annotated with harmonized 32 cell types and 22 molecular tissue regions, as ground truth labels.

FuseMap and other integration or clustering methods were applied to four representative sections. Harmony^12^, scVI^14^, Scanorama^13^, SIMBA^16^, MultiVI^17^, Mario^20^, SpaGCN^21^, GraphST^22^, UTAG^23^, STAGATE^24^ were executed using the Python packages ‘harmonypy’ (v0.0.9), ‘scvi-tools’ (v0.20.3), ‘scanorama’ (v1.7.3), ‘simba’ (v1.2), ‘scvi-tools’ (v0.20.3), ‘mario’ (August 2023), ‘SpaGCN’ (v1.2.7), ‘GraphST’ (v1.1.1), ‘utag’ (v0.1.1.dev38+g0d3d420), and ‘stagate’ (2022.08.23) respectively. Seurat^15^, StabMap^19^, PRECAST^25^ were executed using the R packages ‘seurat’ (v4.3.0.1), ‘stabmap’ (v0.1.8), and ‘precast’ (v1.6.2), respectively. For each method, we used the default hyperparameter settings and data preprocessing steps as recommended. The generated cell embeddings were visualized by UMAP, colored by sample source or ground truth labels. The performance of cell-type and tissue-region integration on the UMAP was evaluated by standard evaluation metrics^32^ including mean average precision, average silhouette width (ASW) and neighbor consistence for biology conservation evaluation, and entropy of batch mixing, Seurat alignment score, omics layer ASW, and graph connectivity for batch correction evaluation. For biology conservation metrics, we used mean average precision, mean average silhouette width, and mean neighbor conservation. For batch effect removal metrics, we used mean batch entropy, Seurat alignment score, average silhouette width and graph connectivity.

To evaluate the performance of unmeasured gene imputation, we employed a leave-one-gene-out integration and imputation strategy. For a total of 52 overlapped genes across the four sections, we excluded one gene each time from one of four representative sections. Then FuseMap was applied to integrate these modified datasets. Built on the inner product between the single-cell embedding and gene embedding, FuseMap imputed gene expression of all genes. We then calculated Pearson’s *R* between the imputed and measured expression of the excluded gene. We benchmarked on two other approaches. The first approach was to align cells in four sections by dimension reduction and batch effect correction methods including principal component analysis (PCA), Harmony, and UMAP and then calculate the imputed expression of each cell as the average expression of *k* nearest cells (here *k*=50) on UMAP. The second approach was similar but otherwise aligned cells in each section with the scRNA-seq Atlas 8.

### Training on large-scale spatial transcriptomics atlas corpus

We assembled an extensive atlas corpus of recent publicly available mouse brain-wide spatial transcriptomics data, incorporating 18,637,640 cells or spots and 26,665 genes. The atlas corpus includes seven atlases from heterogeneous spatial transcriptomics technological platforms including STARmap PLUS, MERFISH, EEL FISH, Stereo-seq and Slide-seq V2. Atlas 1 from Ref. [1] consists of 1,091,280 cells with 1,022 genes across 17 coronal sections and 3 sagittal sections, from 2 animals. Atlas 2 from Ref. [2] as consists of 3,940,980 cells with 500 genes across 59 coronal sections, from 1 animal. Atlas 3 from Ref. [3] consists of 1,735,308 cells with 1,122 genes across 68 coronal sections, and 6,190,162 cells with 1,147 genes across 150 coronal sections and 25 sagittal sections, altogether from 4 animals. Atlas 4 from Ref. [4] consists of 734,696 cells with 483 genes across 9 brain slices (3 replicates of 3 coronal sections from matched locations with respect to bregma), from ≥1 animal. Atlas 5 from Ref. [5] consists of 127,591 cells and 440 total genes across 1 sagittal section, from 1 animal. Atlas 6 from Ref. [6] consists of 50,140 cells with 25,879 genes across 1 coronal section, from 1 animal. Atlas 7 from Ref. [7] consists of 4,767,483 spots with whole transcriptome across 101 coronal sections, from 1 animal.

### Cell type harmonization in atlas corpus

In original cell-type annotations, Atlas 1 was hierarchically organized into 2 nested levels: 26 main level, and 218 sublevel cell types, and Atlas 2 was hierarchically organized into 4 nested levels: 34 classes, 338 subclasses, 1,201 supertypes and 5,322 clusters of cell types. We leverage the integrated embedding to facilitate the transfer of annotations at hierarchical levels of annotation granularity. Based on the embedding, we first transferred main level A1N or A2N cell annotations to cells across the atlas corpus using a multi-layer perceptron classifier. Then for cells in each main level cell type, we transferred sublevel A1N or A2N cell annotations to cells across the atlas corpus using a multi-layer perceptron classifier. During the classification process, 80% of A1N or A2N cells were used as training sets, 10% cells were hold out as the validation set, and 10% cells were used as test sets. The classification accuracy on the test set was calculated using *sklearn.metrics.accuracy_score*^63^. After annotation transfer, cells across the atlas corpus were hierarchically organized into 26 A1N main level, and 191 A1N sublevel cell types; and into 33 A2N classes, 332 A2N subclasses, 1,177 A2N subclasses, and 4,133 A2N clusters. For cell-type annotation transfer, we applied on Atlas 1 to 6 atlases with single cell resolution. We then identified marker genes of cell types calculated using differentially expressed gene analysis via the *rank_genes_groups* function in Scanpy.

### Tissue region harmonization and refinement across atlas corpus

In original tissue-region annotations, Atlas 1 was hierarchically organized into 2 nested levels: 17 main level, and 106 sublevel molecular tissue regions. We transferred the main level A1N tissue regions to cells across the atlas corpus similar to the approach in cell type annotation transfer. Transfer accuracy was 94.0% on 20% hold out data. Next, to further identify finer brain regions, we applied Leiden clustering^64^ to the spatial embedding of each main level transferred molecular tissue region. Depending on if there is visible region shift on the sublevel, we optionally applied the function *scanpy.external.pp.harmony_integrate* for cell alignment across atlases. We then employed the function *scanpy.tl.leiden* with a resolution from 0.1,0.3,0.5, to 1 to perform joint clustering. We then checked for each cluster if it satisfied the following criteria: distinct spatial separation along with region-specific marker genes, consistent spatial distribution across atlases, and precise alignment with the anatomical CCF. We kept clusters that passed the criteria as identified molecular tissue regions. If the identified molecular tissue regions had high correspondence with previous A1N sublevel tissue region annotation, we named it according to previous nomenclature. The identified molecular tissue regions showed finer structures within previous annotations or new structures beyond previous annotations, we named it according to anatomical nomenclature in the Allen Institute Adult Mouse Atlas. After annotation transfer, cells across the atlas corpus were hierarchically organized into 17 main level, and 146 sublevel molecular tissue regions. Tissue region abbreviations: OB, olfactory bulb; CTX, cerebral cortex; CBX, cerebellar cortex; CNU, cerebral nuclei; TH, thalamus; HY, hypothalamus; MB_P_MY, midbrain, pons, and medulla; FT, fiber tracts; VS, ventricular systems; H, habenula; MYdp, medulla, dorsoposterior part; HPFmo, non-pyramidal area of hippocampal formation; MNG, meninges; ENTm, entorhinal area, medial part; HIP, hippocampal region; DG, dentate gyrus; STR, striatum; CTXpl, cortical plate; LSX, lateral septal complex; PAL, pallidum; HB, hindbrain; CBN, cerebellar nuclei.

### Pipeline to build the molecular brain CCF (molCCF)

After different atlases were integrated and annotated with harmonized sublevel tissue region annotations, we registered tissue sections to the anaCCFv3.

#### Order 2D tissue sections

We ordered 405 coronal tissue sections along the anterior to posterior axis and 29 sagittal tissue sections along the lateral to medial axis. Specifically, the 405 coronal tissue sections include 17 from Atlas 1, 59 from Atlas 2, 68 from Atlas 3 replicate 1, 150 from Atlas 3 replicate 2, 9 from Atlas 4, 1 from Atlas 6, and 101 from Atlas 7. The 29 sagittal tissue sections include 3 from Atlas 1, 25 from Atlas 3, and 1 from Atlas 5. Individually, coronal or sagittal tissue sections within each atlas are sorted according to the experimental sectioning locations. To collectively sort coronal or sagittal tissue sections across atlases, we computed a molecular tissue region composition (MTRC) for each tissue section. The MTRC is a 1 × 146 dimensional vector of number in each of the 146 sublevel molecular tissue regions in one section. For coronal sections sorting, we initially used the 150 sections from Atlas 3 replicate 2 as global tissue pool due to its largest coverage. Then we added the 68 sections from Atlas 3 replicate 1 to the global tissue pool. A MTRC similarity matrix between tissue sections in the global tissue pool and 68 sections from Atlas 3 replicate 1 was calculated by *scipy.spatial.distance.cdist*^65^. The optimal insertion order of the 68 sections from Atlas 3 replicate 1 to the global tissue pool was calculated by finding the most similar tissue sections in the global tissue pool for each new section while the original section order was kept unchanged in the individual atlas. After this optimal matching, the global tissue pool updated with orders of both 150 sections from Atlas 3 replicate 2 and 68 sections from Atlas 3 replicate 1. We then iterated inserting coronal tissue sections from other atlases to the global tissue pool and manually adjust the order when necessary. Sagittal tissue sections were ordered manually due to its controllable number. In this way, coronal tissue sections were ordered as No. 1, 2, …, 405 and sagittal tissue sections are ordered as No. 1, 2, …, 29.

#### Identify corresponding image template in the anaCCFv3

For each ordered tissue section, we identified the corresponding image template in the anaCCFv3. For example, we set the coronal anaCCFv3 template at one anterior-posterior axis coordinate corresponding to starting coronal section No. 1 and another anterior-posterior axis coordinate corresponding to ending coronal section No. 405. Then we computed pairwise cosine distances of MTRC between adjacent coronal sections and convert it to intermediate anterior-posterior axis coordinates of in-between coronal sections. In this approach, each tissue section was paired with an anaCCFv3 image template.

#### Register to CCFv3

We registered each tissue section (target) to its paired anaCCFv3 image template (template). The registration included two steps of coarse registration and fine registration. The coarse registration aimed to correct major differences in translation, rotation, and scale. Then the fine registration focused on minor misalignments that were not addressed during coarse registration.

We first detected boundary points. Since most target sections are hemi-brain, we registered only the hemi-brain target and template that shares similar brain anatomy. The template was an image where different values represent different anatomical brain regions. We first converted the image into grid coordinates corresponding to the brain regions. Then from grid coordinates, we found the boundary points of the template that delineate the brain contour. This was achieved through convex hull calculation, vertex extraction, and nearest points detection. Similarly, we found boundary points of the target. Registration was calculated between boundary points of the target and template.

For coarse registration, we detected matched anchor points from the target and template. First, we detected anchor points in the target. To start with, we plotted cells in each tissue section as a 2D image where each cell was colored by sublevel molecular tissue regions. Then we manually clicked two points on each plotted image as the midline of each tissue section from dorsal to ventral direction. With the midline, we defined four anchor points based on the following strategy. The first two anchor points were the intersections of the boundary of the tissue section and a line parallel to the midline. The parallel line was drawn to make the distance of intersection maximal. The third and fourth anchor points were the intersections of the boundary of the tissue section and a line vertical to the midline. The vertical line was drawn to make the distance of intersection maximal. Second, we detected anchor points in the template. The midline of each template from dorsal to ventral direction was fixed for each template. We then used the same strategy to determine the four anchor points. Next, we estimated affine transformation using the detected matching anchor points from the target and template. Finally, we applied the calculated transform matrix to the boundary points of the target.

After coarse registration, we refined the transformation of target boundary points to template boundary points using an iterative approach. We calculated the initial distance cost between the transformed target boundary points and the template boundary points. We matched the nearest template boundary point for each target boundary point and estimate the parameters of the affine transformation that best aligns them. We then updated the overall transformation matrix by multiplying the new matrix with the accumulated transformation matrix. This step effectively composed the new transformation with the existing ones. We iterated this process until the distance cost is smaller than 1𝑒 − 5. After the iterative process completes, the accumulated transformation matrix was applied to all cell coordinates in the target tissue section to transform the tissue section into corresponding template.

In this approach we registered 390 coronal sections and 29 sagittal sections with 13,937,638 cells/spots to the molCCF. The cells/spots have 3D global coordinates with harmonized cell annotations within the molCCF.

### Spatial gene imputation

FuseMap imputed gene expression of 26,665 genes for 18,637,640 cells or spots across the atlas corpus. To assess the imputation accuracy, we compared two neighboring tissue sections from different atlases in the 3D molCCF. The two tissue sections were registered to similar anatomical templates, so they had 3D global coordinates. Based on the 3D global coordinates, we binned the cells/spots using a 10 by 10 grid and got aligned grid coordinates between two tissue sections. In this way, we could compare the correlation of averaged gene expression using the aligned grid coordinates.

We computed the imputation correlation in three categories. For example, representative Section 208 from Atlas 1 and Section 209 from Atlas 2 were used as illustration. First, genes measured in Section 209 were divided into shared or distinct genes based on their overlap with the gene panel of Section 208. Then, for shared genes, we calculated correlation between their measured expressions between Section 208 and 209 as Group 1. Similarly, for shared genes, we calculated correlation between imputed expression in Section 208 and measured expressions in Section 209 as Group 2. For distinct genes, we calculate correlation between their measured expressions between Section 208 and 209 as Group 3. Pearson’s *R* is calculated using *scipy.stats.pearsonr*.

### Gene-to-gene relationship on the learned gene embedding

We utilized a PANTHER dataset with classification of complete protein-coding gene families in mouse genomes and check the distribution of genes from same protein family or gene ontology category on the learned gene embedding UMAP. For genes within each protein family, we computed pairwise distance. Meanwhile we randomly sampled an equal number of genes nonrepetitively from the whole gene set 1,000 times and computed the pairwise distance between randomly selected genes. We performed statistical tests between distances of genes within the same protein family with randomly distributed genes by calculating a Mann-Whitney-Wilcoxon test with *statannotations.Annotator*. We found that statistically genes from the same group tended to be closer on the UMAP. We also calculated a less distribution Mann-Whitney-Wilcoxon test with *scipy.stats.mannwhitneyu* and applied Benjamini-Hochberg correction for multiple tests with *statsmodels.stats.multitest.multipletests*. In this way, we identified 199 protein families with significant closeness.

### Selection of targeted gene panel

#### Task design

The task is to identify a small number of highly informative target genes (i.e., a targeted gene panel). Since spatially resolved single-cell mRNA detection measurements for ten thousand of genes are unavailable, we used the scRNA-seq Atlas 8 with 27,998 genes and 160,796 cells to evaluate the selection accuracy. These cells in Atlas 8 are defined as 39 main levels and 134 detailed cell types. We identified a targeted gene panel of 10 to 150 genes and employ the selected gene panels to classify the original transcriptomic cell types. The classification accuracy was calculated with *sklearn.neural_network.MLPClassifier* with *alpha* equals to 1 and *max_iter* equals to 1,000. Each gene panel accuracy was calculated with 5 random seeds.

#### Benchmark selection methods

For benchmark approaches, we used (i) a randomly chosen gene panel and (ii) a standard panel of top highly variable genes through genes ranked in ‘highly_variable_rank’ value from function *scanpy.pp.highly_variable_genes*.

#### Gene embedding-guided selection method

We first grouped all genes using weighted gene correlation network analysis (WGCNA^66^). This resulted in 13 gene modules. Within each module, we selected *np.floor(#target genes/13)* genes of top highly variable genes and combine selected subpanels together as the final gene panel. The process ensured an even distribution of genes across all 13 gene modules.

### Cell-cell interactions analysis

We followed the previous cell–cell interaction analysis^3^. First, we used harmonized 191 A1N cell subtypes and 17 main level molecular brain tissue regions. We used single cells registered to the molCCF with the molCCF coordinates. Within each region, we calculated the percentage of cells in each subtype and only kept subtypes within sufficient abundance for the following cell–cell interaction analysis. We used a distance threshold of 30 μm, considering that some cell subtype pairs that communicate through paracrine signaling, to compute the number of contact or proximity cell pairs. We then compared against a null distribution created by 1,000 random spatial shifts of cells within a 100 μm radius. Subtype pairs showing significant probability of interaction, determined by adjusted *P* values (<0.05) and observed proximal pair count (>50), were identified as interacting cell-type pairs. We performed the analysis on cells in Atlas 1, 2, and 3 individually and all single cells in the molCCF.

Next, we discerned detained cell-cell interactions using 146 sublevel molecular brain tissue regions. For repeating interaction pattern of astrocyte subtype AC_5 and non-glutamatergic neuroblast subtype NGNBL_3 in sublevel region CNU_13_1-[SEZ] across Atlas 1, 2 and 3, we performed ligand–receptor analysis. Following Ref. [3], we used the CellChat database^67^ to define the ligand–receptor pairs. The expression levels used here were the imputed gene expression. We compared the expression of ligand-receptor pairs between proximal and non-proximal AC_5 and NGNBL_3 cell pairs. Ligand-receptor expression scores were calculated for both types of pairs, with proximal pairs having a soma centroid distance smaller than 30 μm. Statistical tests including Welch’s t-test and the Benjamini–Hochberg correction were used to determine significant differences. Ligand-receptor pairs with high expression in proximal pairs and adjusted p-values below 0.01 were identified as statistically upregulated in proximity, with a threshold distance of 30 μm. Protocadherin pathways were removed due to bias in sequence diversity after manual inspection.

### Benchmark annotation transfer performance

We used the four representative sections in the method description of ‘Benchmark integration performance’. We used Section 1 as a base reference, and combined Section 1, 3 and 4 as a large corpus reference. We used Section 2 as query. As previously described, cells in reference and query sections had harmonized cell type labels. We transferred annotations from base reference or large corpus reference to query and evaluate the accuracy of transferred accuracy with ground truth.

FuseMap and other annotation transfer methods were applied. FuseMap mapped query data to the reference embedding through transfer learning. Then based on integrated single-cell embedding, a multi-layer perceptron classifier was trained on reference labeled to predict on query data. Concerto^49^, ScanVI^50^, expiMap^51^, TOSICA^52^, ItClust^53^, MARS^54^, scPoli^55^, and SCALEX^56^ were executed using the Python packages ‘concerto_function5_3(August 2023), ‘scarches’(0.5.9), ‘scarches’(0.5.9), ‘TOSICA’(September 2023), ‘ItClust’(v1.2.0), ‘mars’(August 2023), ‘scarches’(0.5.9), and ‘scalex(v1.0.2) respectively. For each method, we used the default hyperparameter settings and data preprocessing steps as recommended. Evaluation metrics included *accuracy_score*, *precision_score*, *recall_score*, *f1_score*, *normalized_mutual_info_score*, *adjusted_mutual_info_score*, and *adjusted_rand_score* under *sklearn.metrics*.

We also used 17 Atlas 1 and 55 Atlas 2 coronal sections as reference to train a FuseMap model and evaluated label transfer accuracy on hold-out dataset. We evaluated accuracy on four leave-out Atlas 2 sections (‘C57BL6J-638850.05’, ‘C57BL6J-638850.25’, ‘C57BL6J-638850.45’, ‘C57BL6J-638850.66’) spanning in an even interval from anterior to posterior. We also tested on three sagittal sections from Atlas 1 to 4 as reference. We calculated the accuracy between predicted and ground truth cell type or tissue region labels using *sklearn.metrics*.*accuracy_score* and normalized the matrix along the row and column axes sequentially.

### Annotation transfer on various spatial transcriptomics query datasets

We finetuned the pretrained brain FuseMap model on query datasets. First, we studied a STARmap PLUS query dataset with 5,413 genes. Query dataset had originally 12 main level and 39 sublevel cell types and 9 main level anatomical tissue regions. We transferred A1N 22 main level and 141 sublevel cell types, and 11 main level and 81 sublevel molecular tissue regions. Then we studied two MERFISH sections of the mouse primary motor cortex with 256 genes. Query dataset had originally 24 main level and 96 sublevel cell types and no tissue regions available. We transferred A2N 23 classes, 106 subclasses, 186 supertypes, and 6 main level and 22 sublevel molecular tissue regions. Finally, we applied FuseMap to analyze two sections of the mouse olfactory bulb from different chip-capture-based spatial transcriptomics platforms: Slide-seqV2 and Stereo-seq.

### Software

The following packages and software were used in the data analysis^63,65,68-75^. MATLAB R2019b and Python 3.6. ImageJ 1.51, R 4.0.4, RStudio 1.4.1106, Jupyter Notebook 6.0.3, Anaconda 2-2-.02, h5py 3.1.0, hdf5 1.10.4, matplotlib 3.1.3, seaborn 0.11.0, scanpy 1.6.0, numpy 1.19.4, scipy 1.6.3, pandas 1.2.3, scikit-learn 0.22, umap-learn0.4.3, pip 21.0.1, numba 0.51.2, tifffile 2020.10.1, scikit-image 0.18.1, anndata 0.8.0, itertools 8.0.0, diffxpy 0.7.4, and gprofiler2 package v0.1.9. Other detailed information of benchmark packages and software are in the ‘Benchmark FuseMap performance’ and ‘Benchmark annotation transfer performance’ section.

### Statistical analysis

Pearson’s *R* and its *P* values (two-tailed) in Fig. 2b, Fig. 4c-h, and Extended Data Fig. 9c,d were calculated with *scipy.stats.pearsonr. P* values in Fig. 4j were calculated with Mann-Whitney-Wilcoxon test with *statannotations.Annotator*. *P* values in Extended Data Fig. 10 were calculated following the previous package^3^ with *scipy.stats.norm.sf* of *z* scores and then corrected with Benjamini-Hochberg correction for multiple tests with *statsmodels.stats.multitest.multipletests*.

### Data Availability

Publicly available datasets are used in this study (summarized in Supplementary Table 1). The spatial transcriptomics atlases of mouse brain are available in https://singlecell.broadinstitute.org/single_cell/study/SCP1830 (Atlas 1), https://doi.brainimagelibrary.org/doi/10.35077/g.610 (Atlas 2), https://doi.brainimagelibrary.org/doi/10.35077/act-bag (Atlas 3), https://info.vizgen.com/mouse-brain-map (Atlas 4), http://mousebrain.org/adult/ (Atlas 5), https://db.cngb.org/search/project/CNP0001543/ (Atlas 6), https://www.braincelldata.org/ (Atlas 7). The PANTHER classification of complete protein-coding gene families in mouse genomes is available in http://data.pantherdb.org/ftp/sequence_classifications/current_release/PANTHER_Sequence_Classificatio <u>n_files/PTHR18.0_mouse</u>. The scRNA-seq dataset is available in https://mousebrain.org/. The STARmap PLUS query dataset is available in https://zenodo.org/records/8041114. The MERFISH query dataset is available in https://doi.brainimagelibrary.org/doi/10.35077/g.21. The Stereo-seq query dataset is available in https://github.com/JinmiaoChenLab/SEDR_analyses/tree/master/data. The Slide-seqV2 data can be downloaded from https://singlecell.broadinstitute.org/single_cell/study/SCP815/.

### Code Availability

The source code is available on GitHub at https://github.com/wanglab-broad/FuseMap and https://github.com/LiuLab-Bioelectronics-Harvard/FuseMap. We also provide an interactive online database (http://fusemap.spatial-atlas.net/) for exploratory analysis of the molCCF. All original code will be deposited at GitHub and publicly available as of the date of publication.

## Acknowledgements

We thank Z. Lin, W. Wang and G. Dong for insightful discussions. X.W. acknowledges support from Stanley Center for Psychiatric Research, Edward Scolnick Professorship, Ono Pharma Breakthrough Science Initiative Award, Packard Fellowship for Science and Engineering, Sloan Research Fellowship, and NIH DP2 New Innovator Award and Brain Initiative. J.L. acknowledges support from the Aramont Fund. W.X.W. is a Damon Runyon (National Mah Johngg League, Inc) Fellow, supported by the Damon Runyon Cancer Research Foundation (DRG-#2512-23).

## Author contributions

X.W., J.L., and Y.H. conceived the idea. Y.H. developed the framework, designed computational and data analyses, prepared figures, and drafted the manuscript. H.Sheng contributed critical discussions and input on the figures. H.Shi and W.X.W. provided input on biological interpretations and edits during revision. Z.T. set up the online data portal. Y.H., J.L., and X.W. revised the manuscript with input from all authors. X.W. and J.L. supervised the study.

## Additional information

Correspondence and requests for materials should be addressed to Xiao Wang or Jia Liu.

## Supplementary information

**Extended Data Figure 1.**
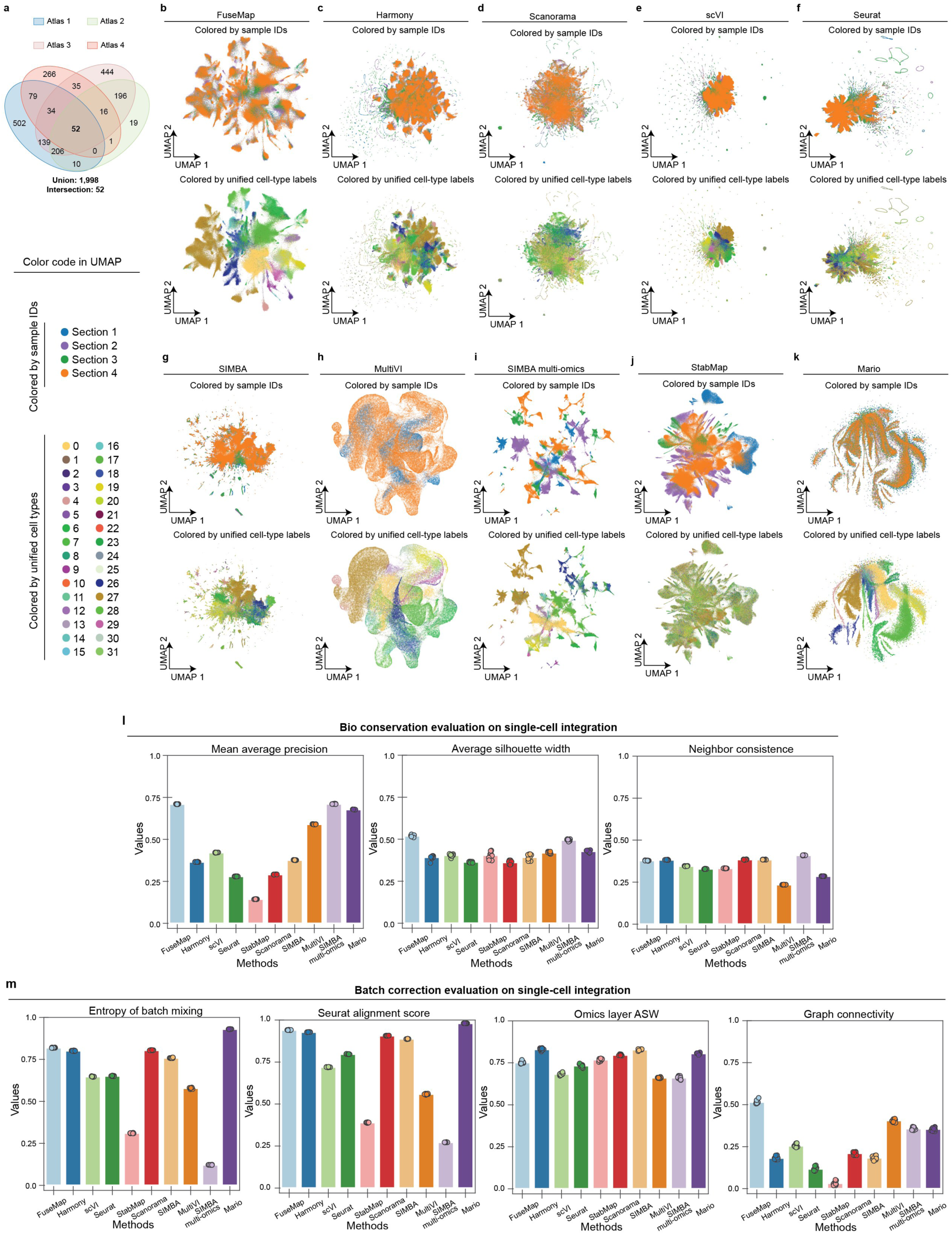
Evaluation of cell-type integration performance. **a**, Intersection size of gene panels in the four representative sections. **b-k**, UMAP of cell-type embedding from FuseMap and other methods, colored by sample IDs (upper row) and ground truth labels (lower row). **l**,**m**, Accuracy of cell-type integration methods on bio conservation (**k**) and batch correction (**l**) evaluation metrics.

**Extended Data Figure 2.**
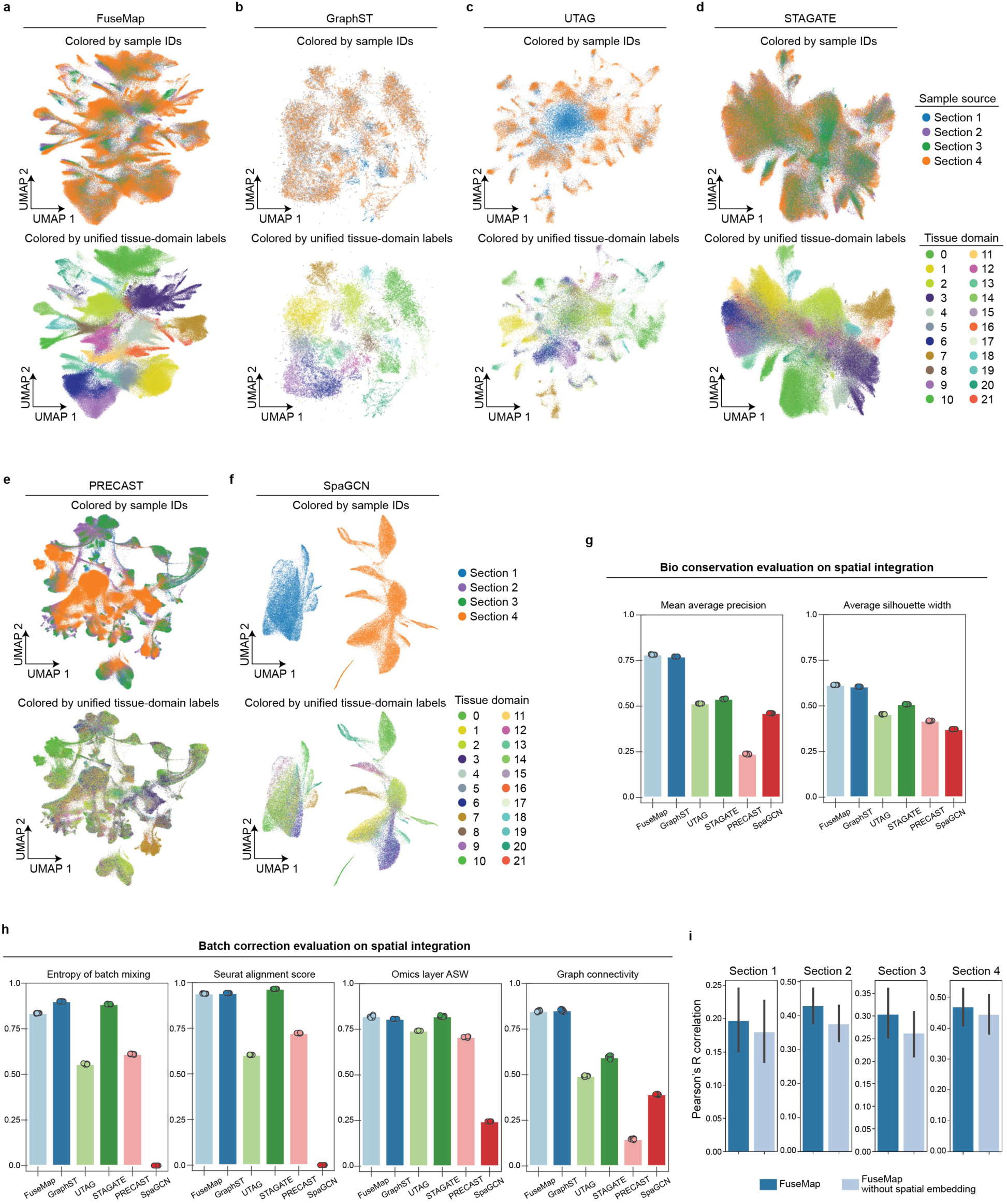
Evaluation of tissue-region integration performance. **a-f**, UMAP of tissue-region embedding from FuseMap and other methods, colored by sample IDs (upper row) and ground truth labels (lower row). **g**,**h**, Accuracy of cell-type integration methods on bio conservation (**g**) and batch correction (**h**) evaluation metrics. **i**, Comparison of gene imputation performance between FuseMap with and without spatial embedding on four representative sections across platforms.

**Extended Data Figure 3.**
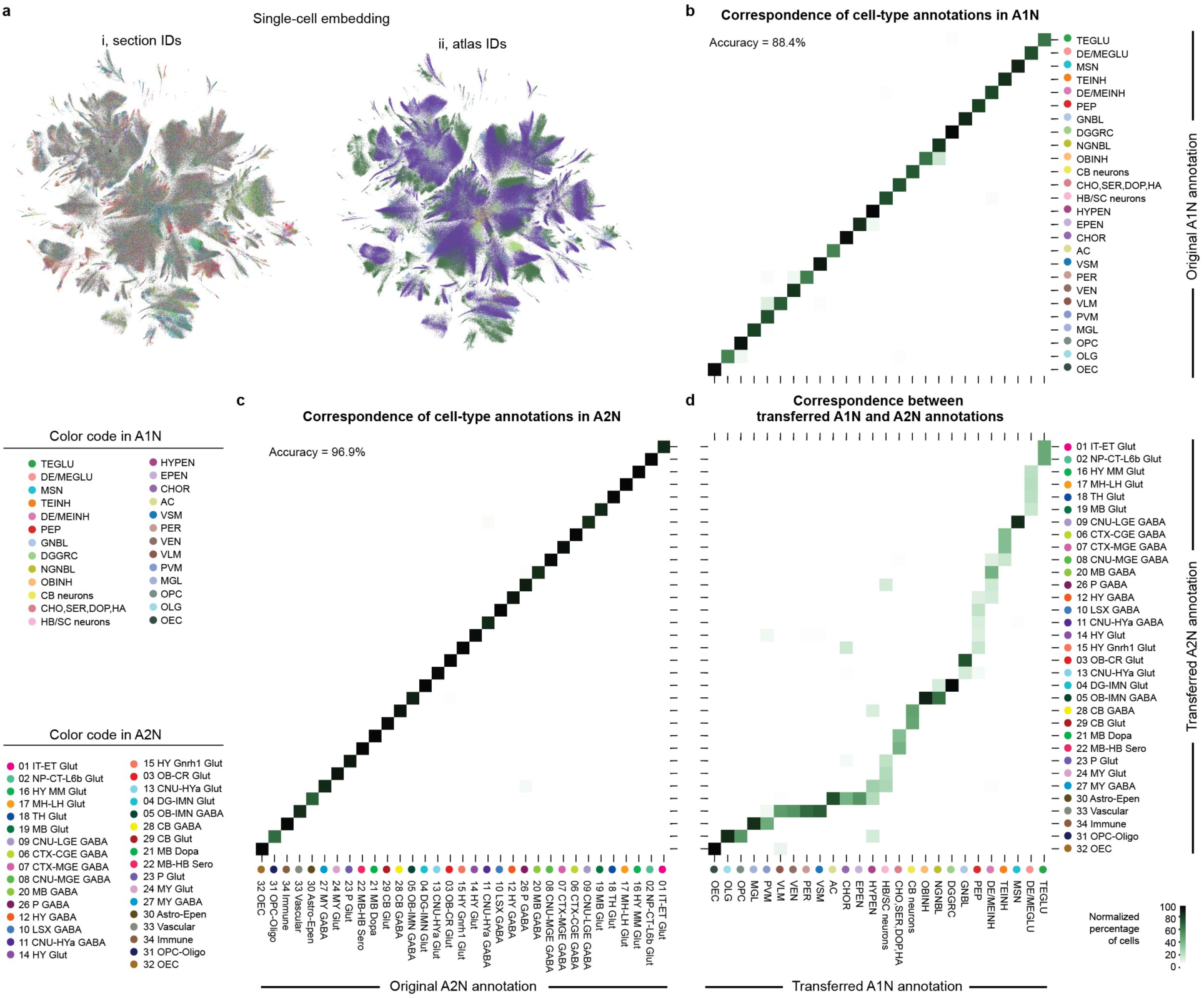
Harmonization of cell types in the atlas corpus. **a**, UMAP of cell-type embedding of randomly sampled one million cells in the atlas corpus, colored by section IDs (i) and atlas IDs (ii). **b**, Percentage of original (rows) within transferred (columns) cell-type annotations from A1N. **c**, Percentage of original (columns) within transferred (rows) cell-type annotations from A2N. **d**, Percentage of transferred cell-type annotation from A1N (columns) within transferred cell-type annotation from A2N (rows).

**Extended Data Figure 4.**
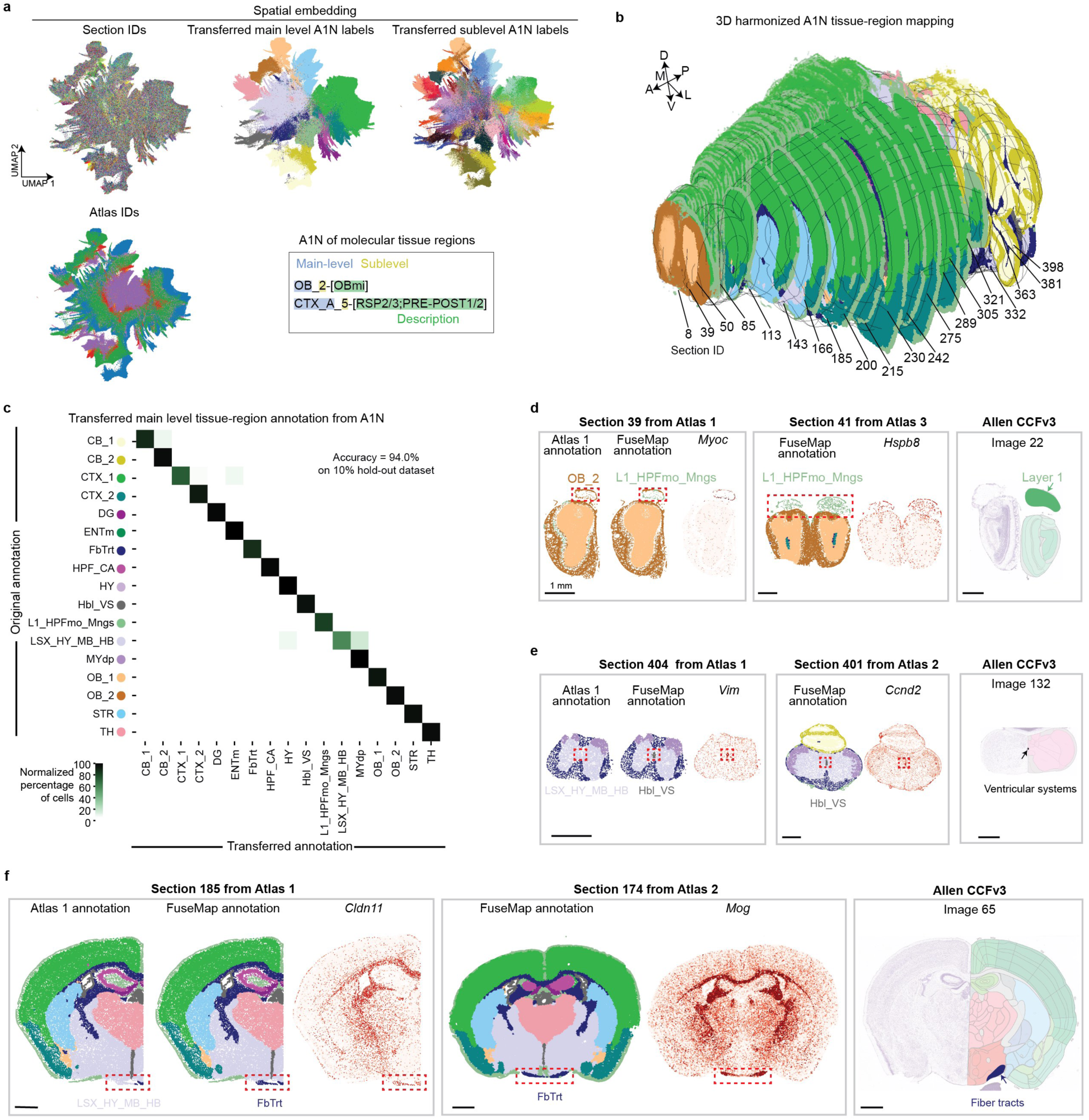
Harmonization of main level tissue regions in the atlas corpus. **a**, UMAP of tissue-region embedding of randomly sampled one million cells in the atlas corpus, colored by section IDs, atlas IDs, transferred main and sub annotation from Atlas 1 nomenclature (A1N). **b**, 3D harmonized tissue-region mapping of 434 sections (distal hemisphere), with example coronal tissue sections highlighted (proximal hemisphere) in the molCCF. **c**, Percentage of original tissue-region annotation (rows) within transferred tissue-region annotation (columns) from A1N. **d-f**, Representative main level tissue region refinements in Section 39, Section 404, and Section 185, illustrated with distribution in neighboring tissue sections and marker gene expression. Scale bar, 1 mm.

**Extended Data Figure 5.**
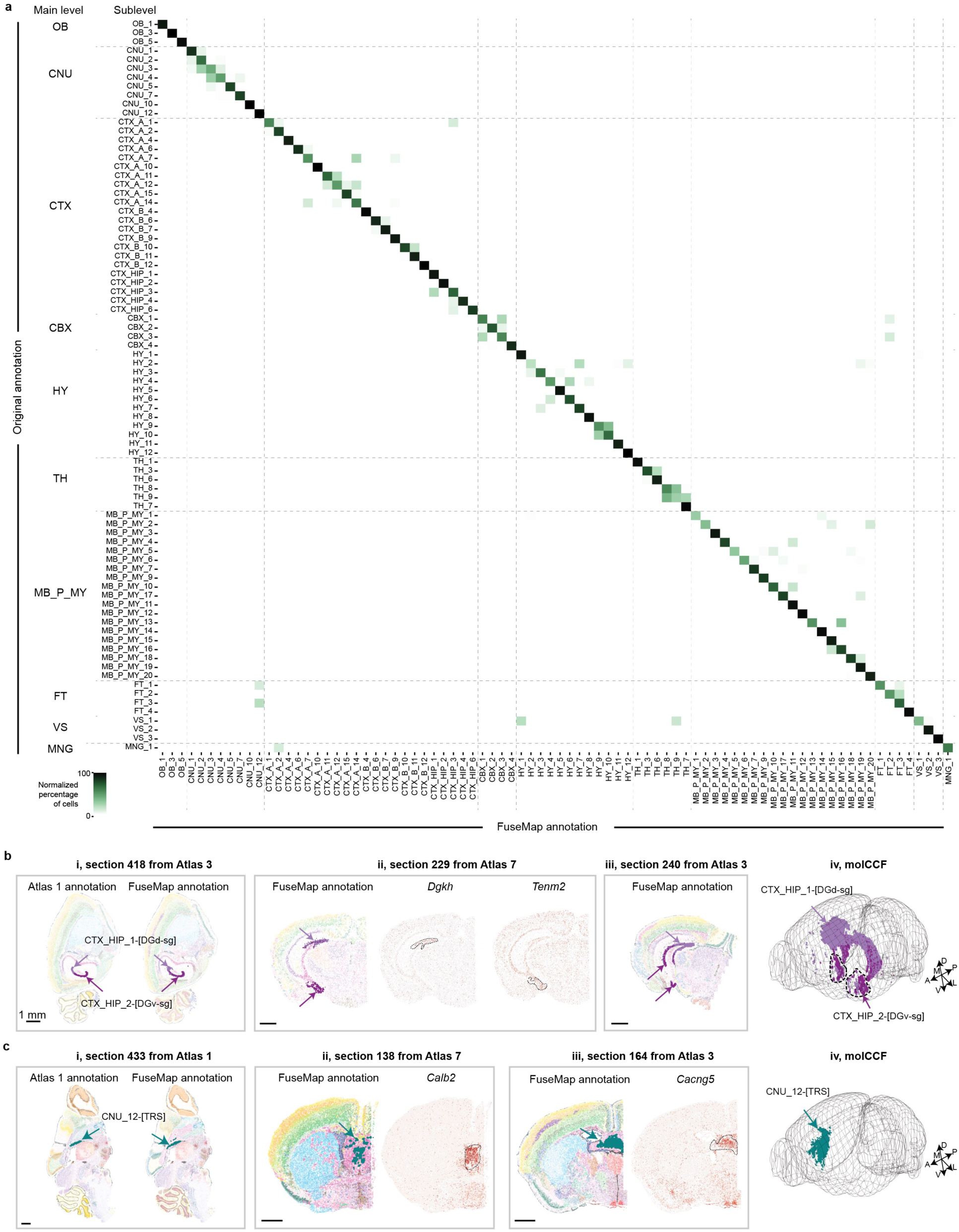
Harmonization of sublevel tissue regions across brain atlases. **a**, Percentage of original A1N (rows) with transferred (columns) molecular tissue region annotations at sublevel. **b**,**c**, Representative sublevel tissue regions in CTX_HIP_1 and CTX_HIP_2 (**b**), and in CNU_12 (**c**), illustrated with distribution in representative tissue sections and the molCCF along with marker gene patterns. DGd-dg, dentate gyrus, dorsal part, granule cell layer; DGv-dg, dentate gyrus, ventral part, granule cell layer; TRS, triangular nucleus of septum. Scale bar, 1 mm.

**Extended Data Figure 6.**
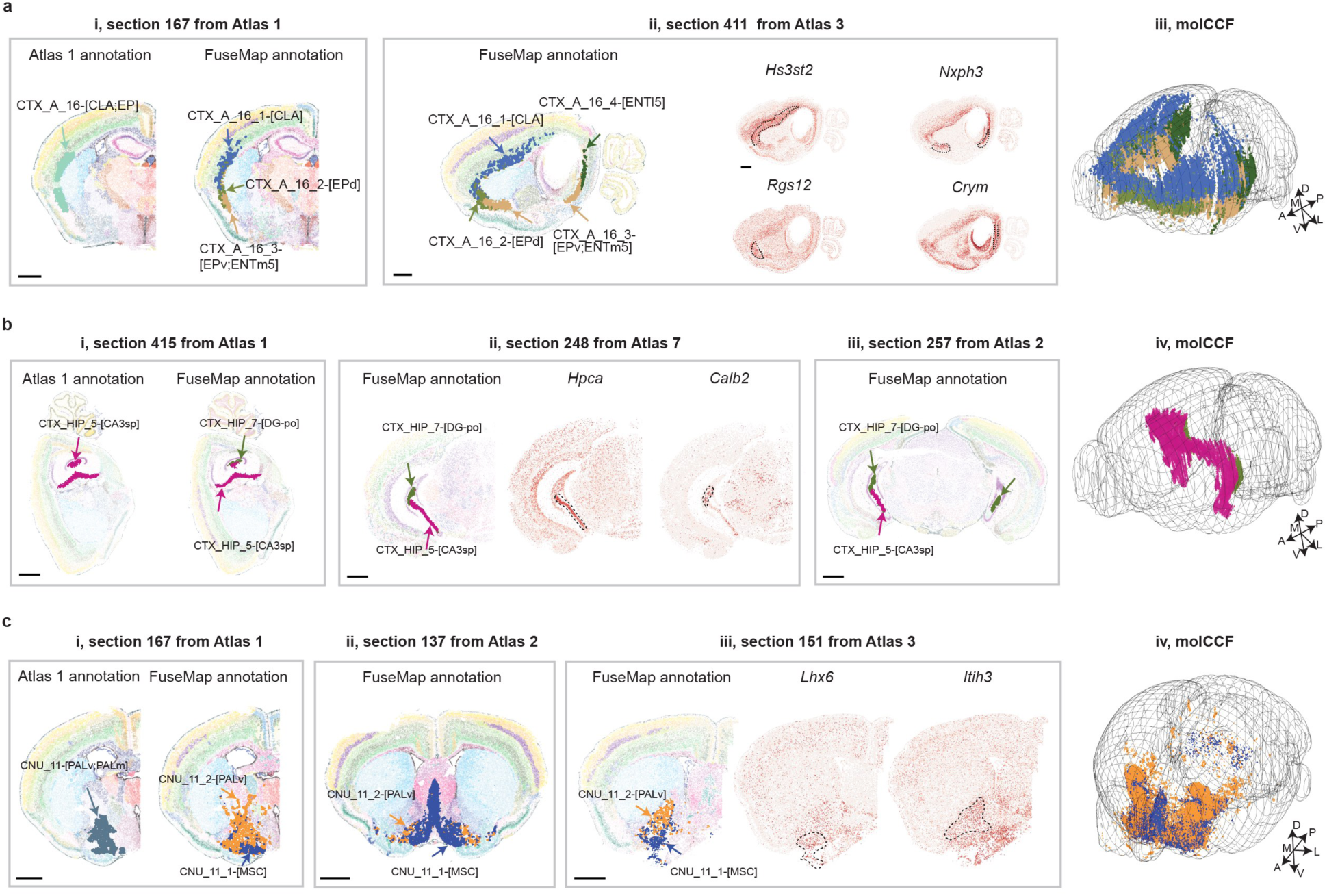
Additional examples of newly identified molecular tissue regions across the mouse brain. **a-e**, Representative newly identified sublevel tissue regions in A1N CTX_A_16 (**a**), A1N CTX_HIP_5 (**b**), and A1N CNU_11 (**c**), illustrated with distribution in representative tissue sections and the molCCF along with marker gene patterns. CLA, claustrum; EP, endopiriform nucleus; EPd, EP dorsal part; EPv, EP ventral part; ENTm, entorhinal area, medial part, dorsal zone; ENTl, entorhinal area, lateral part; CA3sp, field CA3, pyramidal layer; DG-po, dentate gyrus, polymorph layer; PALv, pallidum, ventral region; PALm, pallidum, medial region; MSC, medial septal complex. Scale bar, 1 mm.

**Extended Data Figure 7.**
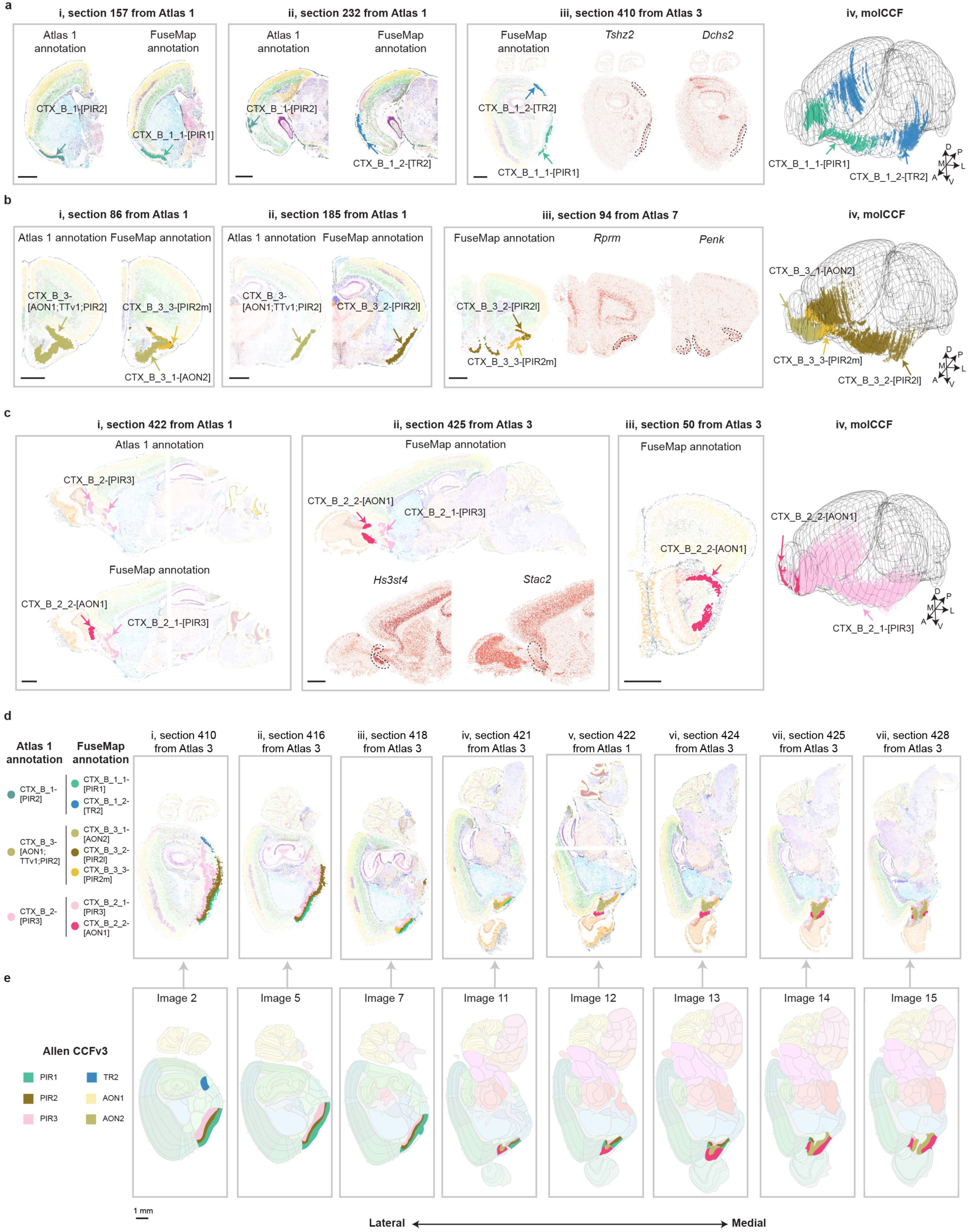
Examination of molecular piriform areas. **a-c**, Representative newly identified sublevel tissue regions in A1N CTX_B_1 (**a**), A1N CTX_B_3 (**b**), A1N CTX_B_2 (**c**), illustrated with distribution in representative tissue sections and the molCCF along with marker gene patterns. PIR, piriform area; PIR1, PIR molecular layer; PIR2, PIR pyramidal layer; PIR3, PIR polymorph layer; AON, anterior olfactory nucleus; TR, postpiriform transition area. **d**,**e**, Spatial distribution of PIR-related molecular tissue region in representative sections across atlases (**d**) and anatomical tissue maps in the anaCCFv3 (**e**) along the lateral–medial axis. Scale bar, 1 mm.

**Extended Data Figure 8.**
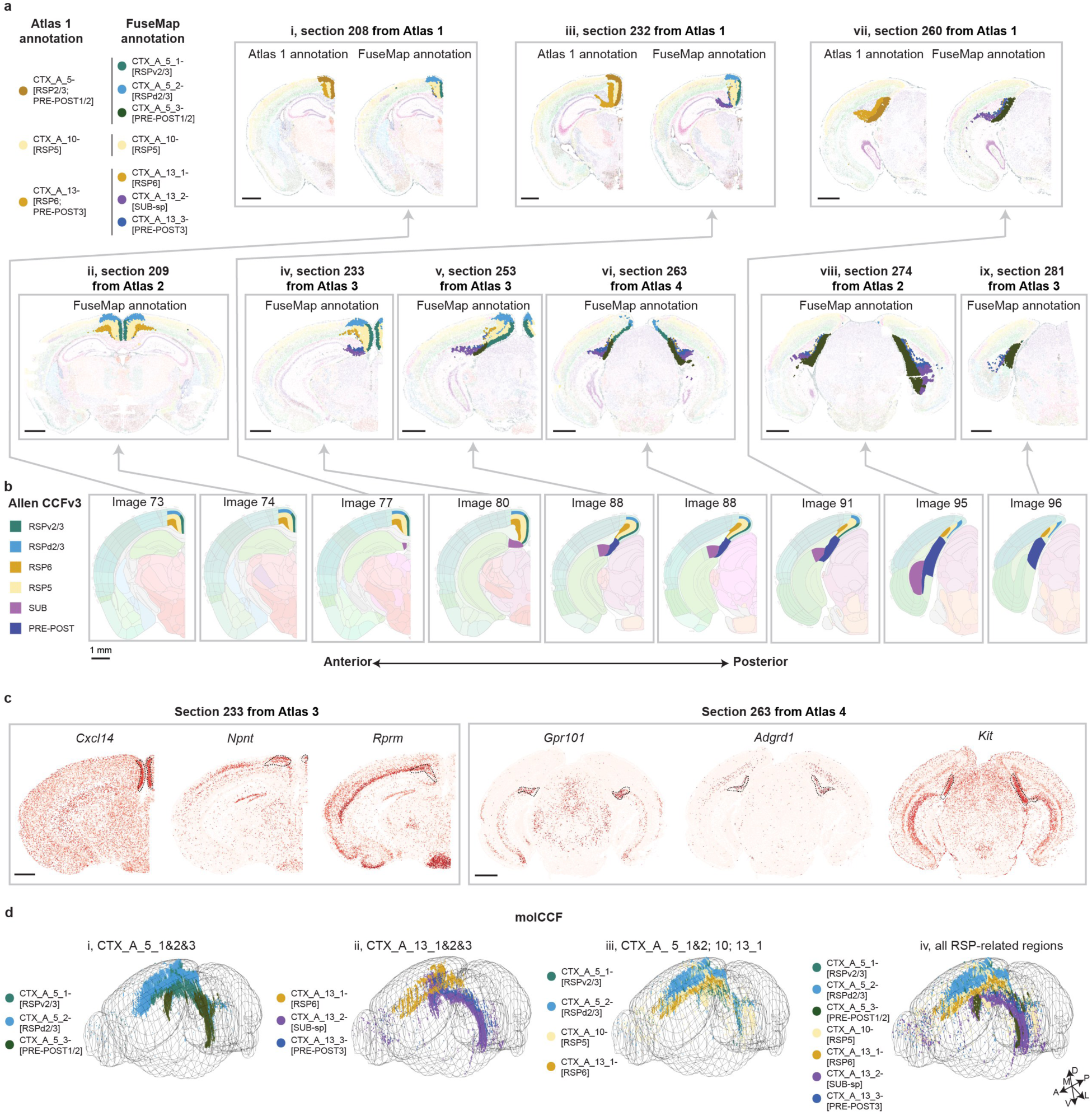
Examination of molecular retrosplenial areas. **a,b**, Spatial distribution of RSP-related molecular tissue region in representative sections in Atlas 1 (**a**, upper row) and other atlases (**b**, lower row), and anatomical tissue maps in the anaCCFv3 (**b**) along the anterior– posterior axis. RSP, retrosplenial area; POST, postsubiculum; PRE, presubiculum; SUB, subiculum. **c**, Spatial expression patterns of marker genes in RSP-related molecular tissue regions. **d**, 3D distribution of RSP-related molecular tissue regions in the molCCF. Scale bar, 1 mm.

**Extended Data Figure 9.**
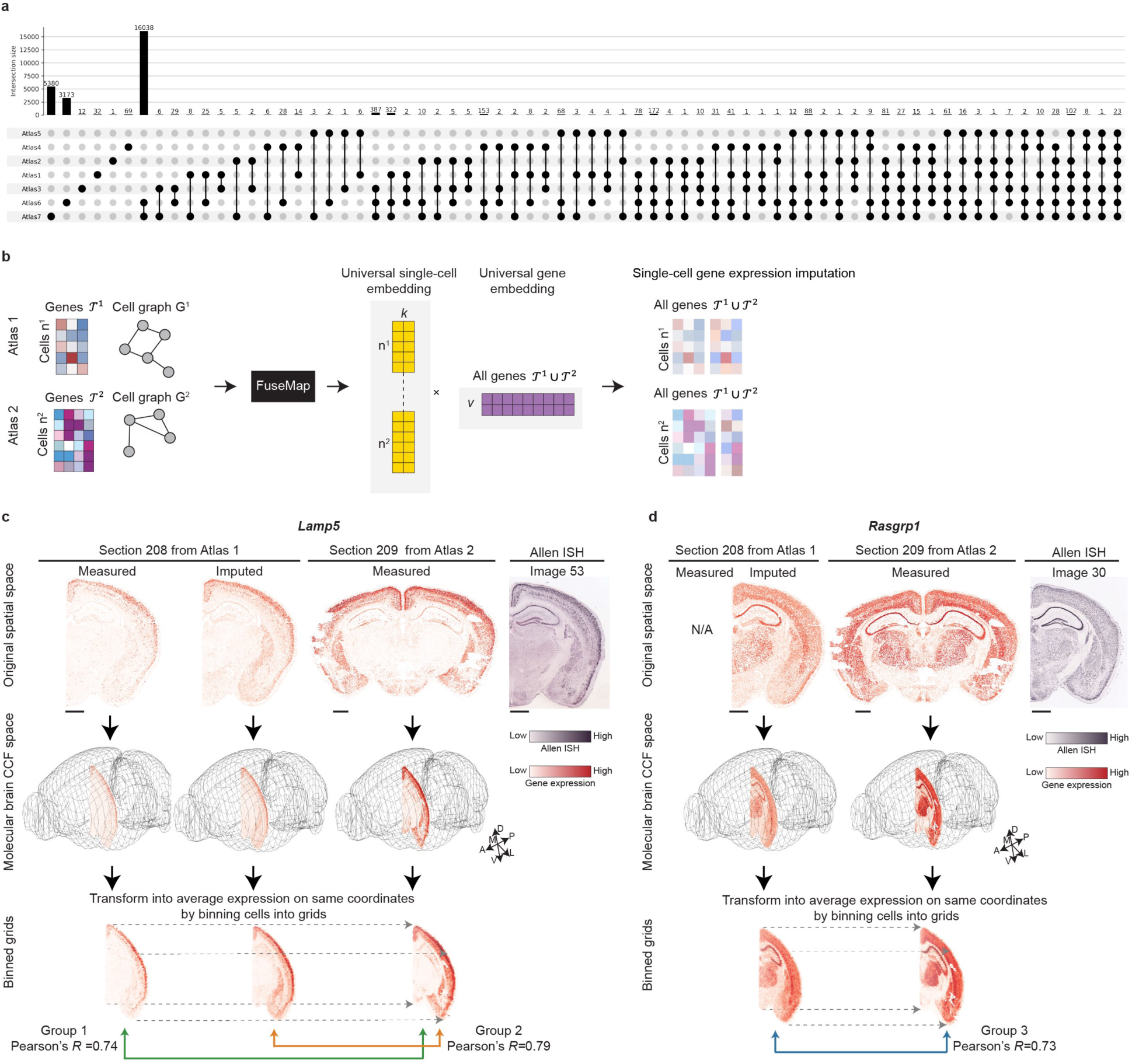
Evaluation of single-cell gene expression imputation. **a**, Intersection size of gene panels in Atlases 1 to 7. **b**, Schematics of single-cell gene imputation method. **c**,**d**, Workflow to compute imputation accuracy. Two neighboring tissue sections from the 3D molCCF are compared. Cells or spots in the two sections are binned into identical coordinates based on their alignment within the molCCF. Correlations are calculated between measured expressions of a shared gene *Lamp5* (**c**, Group 1), imputed versus measured expressions for shared genes *Lamp5* (**c**, Group 2), and imputed versus measured expressions for distinct genes *Rasgrp1* (**d**, Group 3). Scale bar, 1 mm.

**Extended Data Figure 10.**
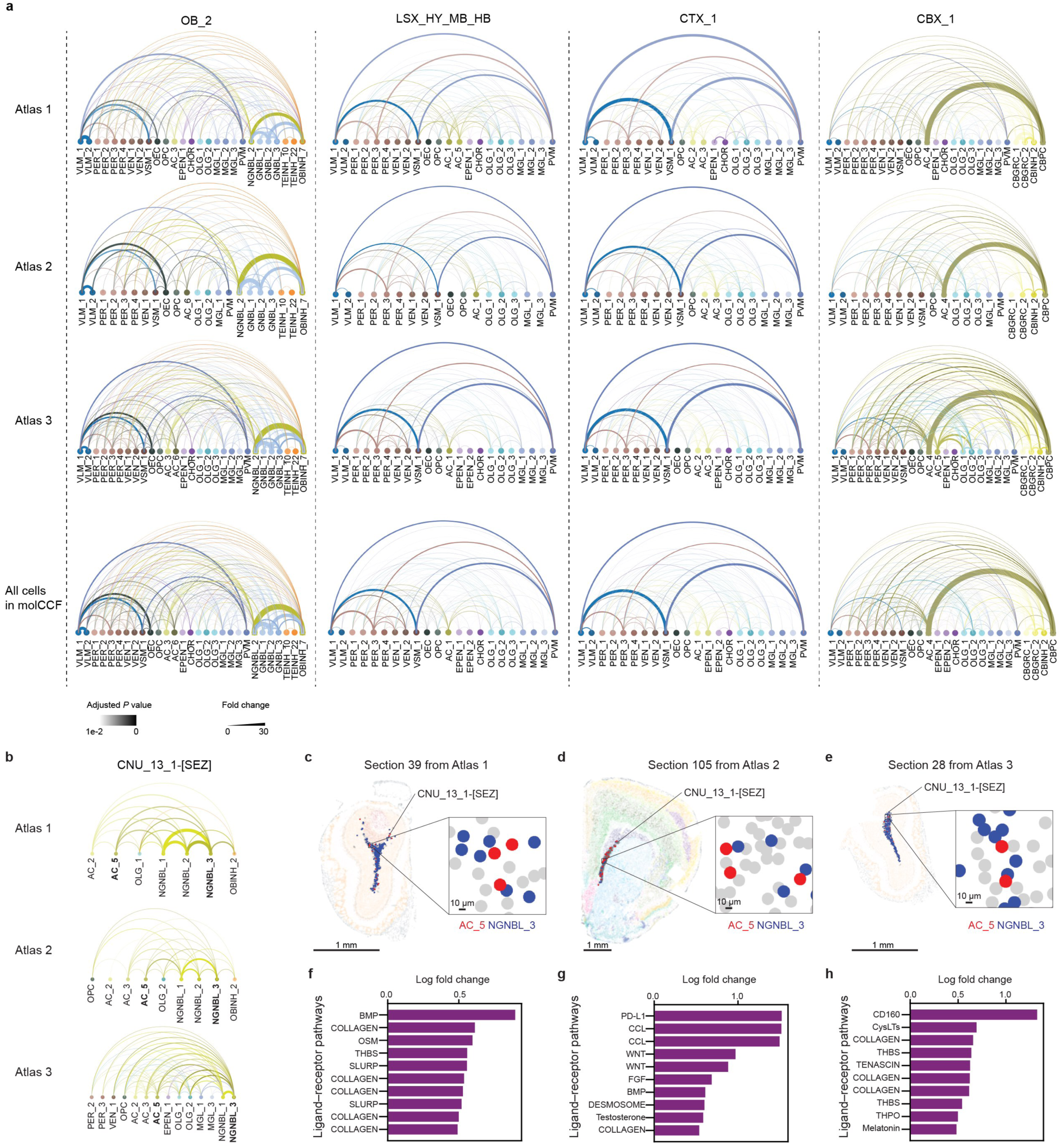
Cell-cell interactions across atlases. **a**, Main level molecular brain-region-specific cell-cell adjacency in Atlases 1, 2, and 3, respectively. Representative regions of OB_2, LSX_HY_MB_HB, CTX_1, and CBX_1 are shown from left to right. **b**, Sublevel molecular brain region CNU_13_1-[SEZ] specific cell-cell adjacency in Atlases 1, 2, and 3 respectively. Each line corresponds to a cell-type pair, with the linewidth indicating fold change in proximity frequency compared with random chance and transparency indicating *P* values corrected by the Benjamini–Hochberg procedure. **c**,**d**,**e**, Representative brain sections from Atlases 1, 2 and 3, respectively. AC_5 and NGNBL_3 cells in CNU_13_1-[SEZ] are colored in red or blue. **f**,**g**,**h**, Ligand–receptor pathways upregulated in the AC_5 and NGNBL_3 proximal cell pairs as compared to non-proximal cell pairs from Atlases 1, 2 and 3, respectively. FbTrt, fiber tracts; LSX_HY_MB_HB, Lateral septal complex/hypothalamus/midbrain/hindbrain; STR, striatum; CTX_1, Cerebral cortex, dorsal part; OB_2, Olfactory bulb outer layers (glomerular, nerve layer, etc); CBX_1, Cerebellar granule layer and Purkinje cell layer. AC, astrocytes; NGNBL, non-glutamatergic neuroblasts. Scale bar, 1 mm.

**Supplementary Figure 1.**
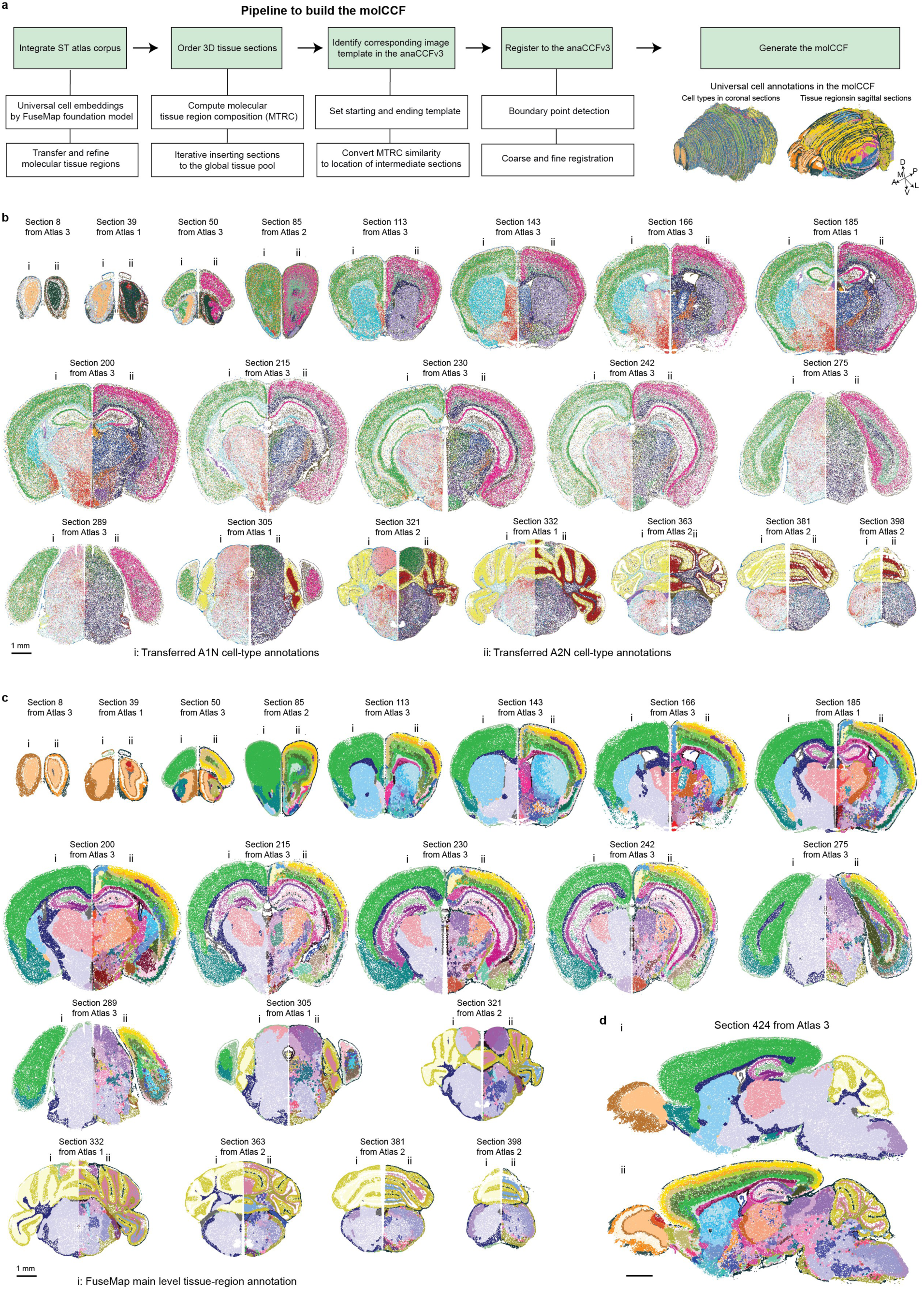
Workflow to build the molCCF. **a**, Pipeline to build the molCCF. **b**, Example coronal tissue sections highlighted in Fig. 2d showing transferred cell types from A1N (i) and Atlas 2 (ii). **c**,**d**, Example coronal (**c**) and sagittal (**d**) sections showing main level (i) and sublevel (ii) transferred molecular tissue regions. Scale bar, 1 mm.

**Supplementary Figure 2.**
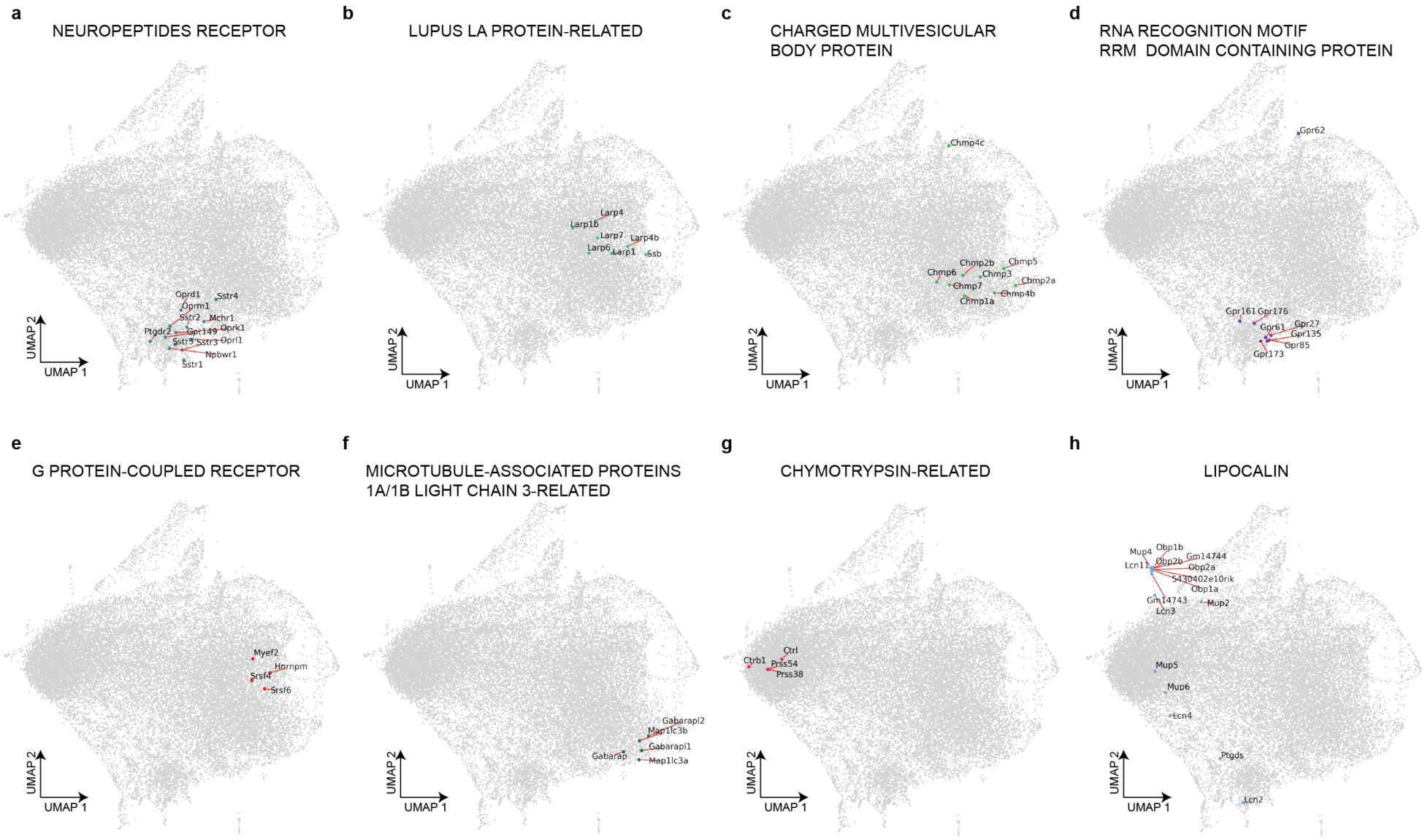
Example of PANTHER protein-coding gene families on UMAP of the gene embedding. **a-h**, UMAP of the gene embedding with gene symbols from one PANTHER protein-coding gene family labeled.

**Supplementary Figure 3.**
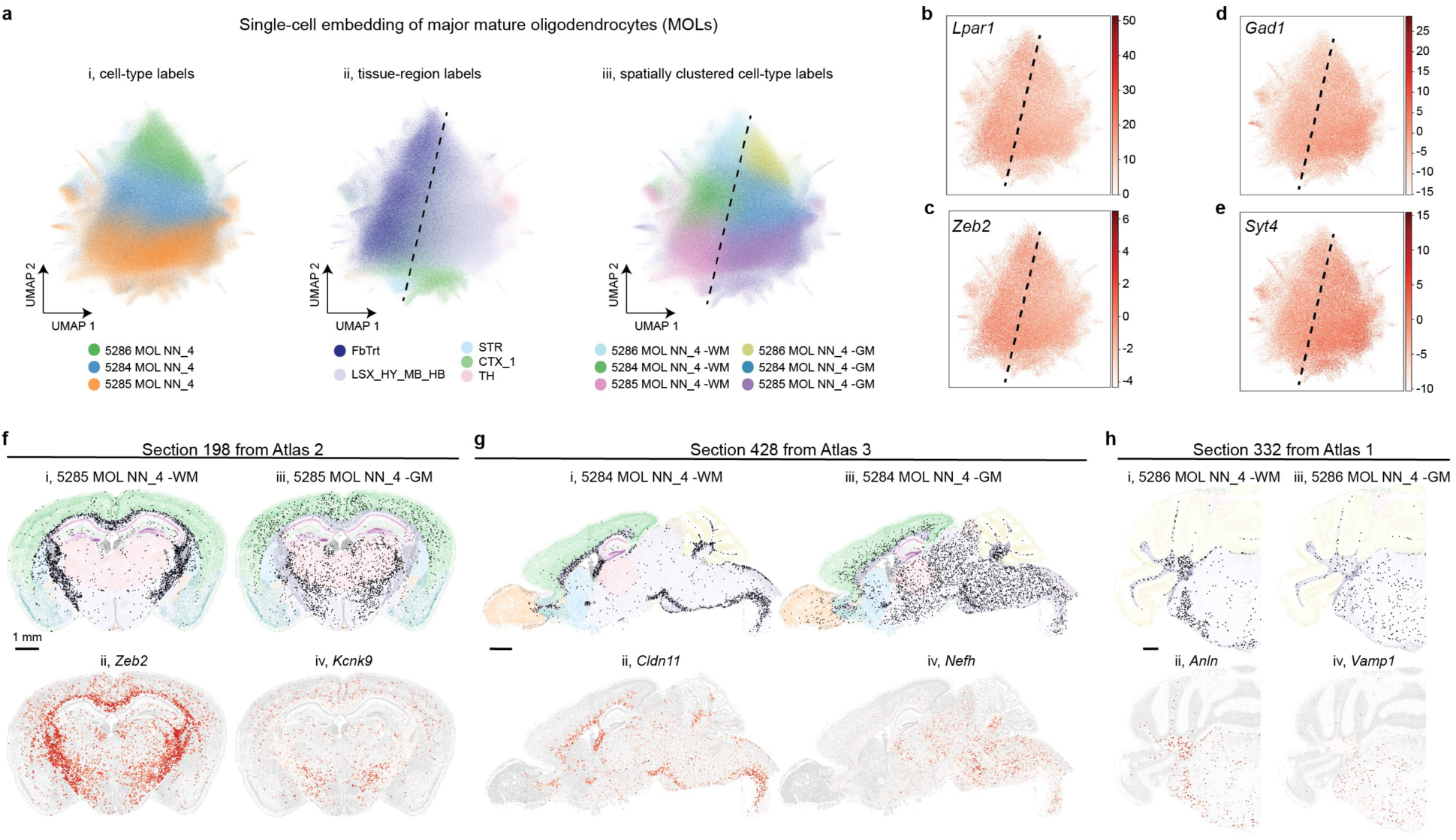
Spatially informed clustering of mature oligodendrocytes (MOLs). **a**, UMAP of the single-cell embedding of three major MOLs, colored by (i) cell-type labels, (ii) tissue-region labels, and (iii) spatially clustered cell-type labels. **b-e**, UMAP of the single-cell embedding of *Lpar1*, *Zeb2*, *Gad1*, and *Syt4*. The expression levels used here are the imputed gene expression. **f**, Distribution of spatially clustered cell subtype 5285 MOL NN_4 -WM (i) and 5285 MOL NN_4 -GM (iii), along with corresponding marker gene *Zeb2* (ii) and *Kcnk9* (iv) on representative section 198 from Atlas 2. The expression levels used here are the measured gene expression. **g**, Distribution of spatially clustered cell subtype 5284 MOL NN_4 -WM (i) and 5284 MOL NN_4 -GM (iii), along with corresponding marker gene *Gfap* (ii) and *Nefh* (iv) on representative section 428 from Atlas 3. **h**, Distribution of spatially clustered cell subtype 5286 MOL NN_4 -WM (i) and 5286 MOL NN_4 -GM (iii), along with corresponding marker gene *Anln* (ii) and *Vamp1* (iv) on representative section 332 from Atlas 1. Scale bar, 1 mm.

**Supplementary Figure 4.**
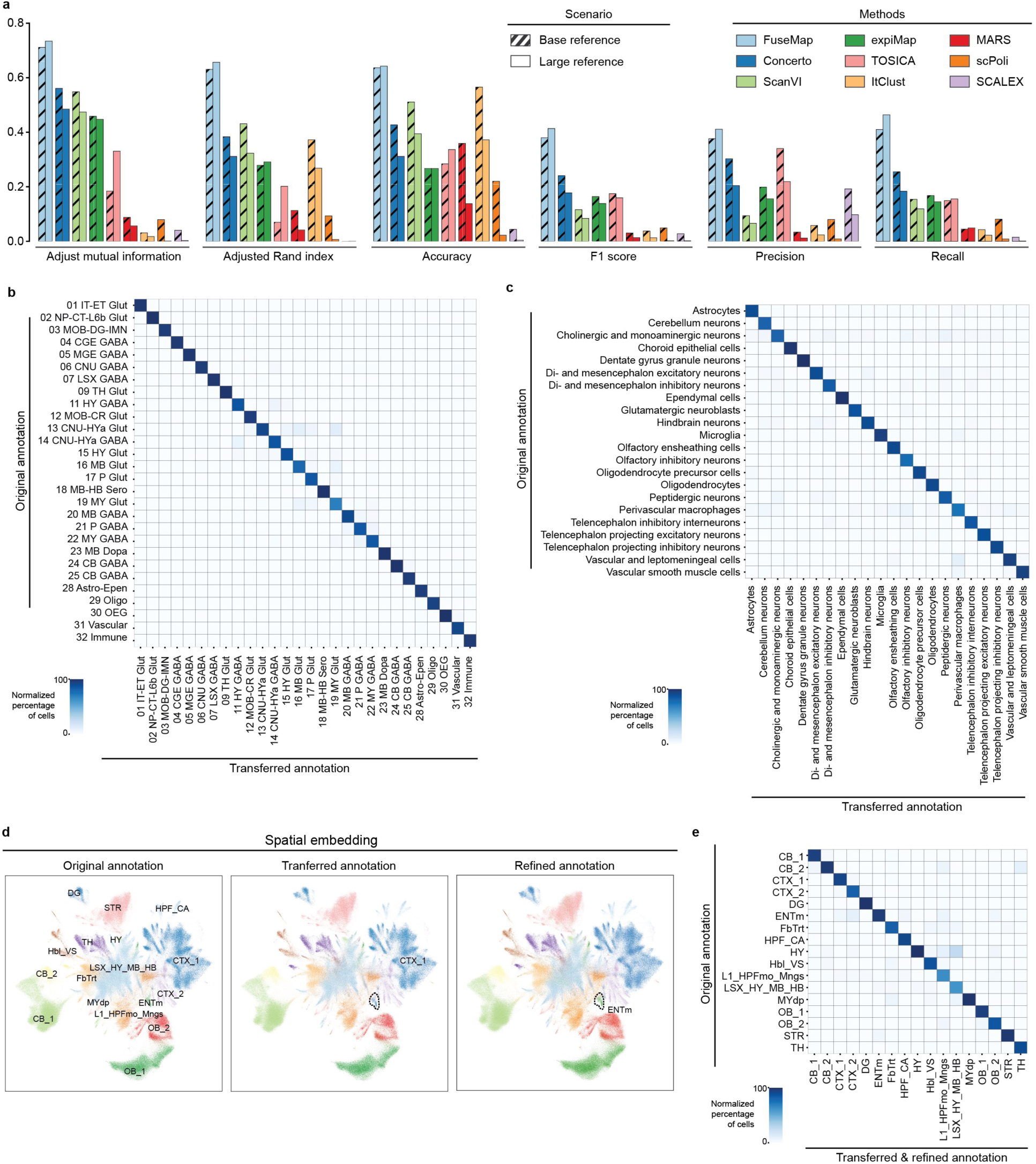
Benchmark performance on cell-type annotation transfer task. **a**, Benchmark cell-type annotation transfer performance on representative sections from Atlases 1 to 4. **b**, Percentage of original (rows) with transferred (columns) cell type annotations on hold-out Atlas 3 sections. **c**, Percentage of original (rows) with transferred (columns) cell type annotations on hold-out Atlas 1 sections. **d**, UMAP of spatial embedding colored by original, transferred, and refined tissue region annotations on hold-out Atlas 1 sections. **e**, Percentage of original (rows) with transferred and refined (columns) tissue region annotations on hold-out Atlas 1 sections.

**Supplementary Figure 5.**
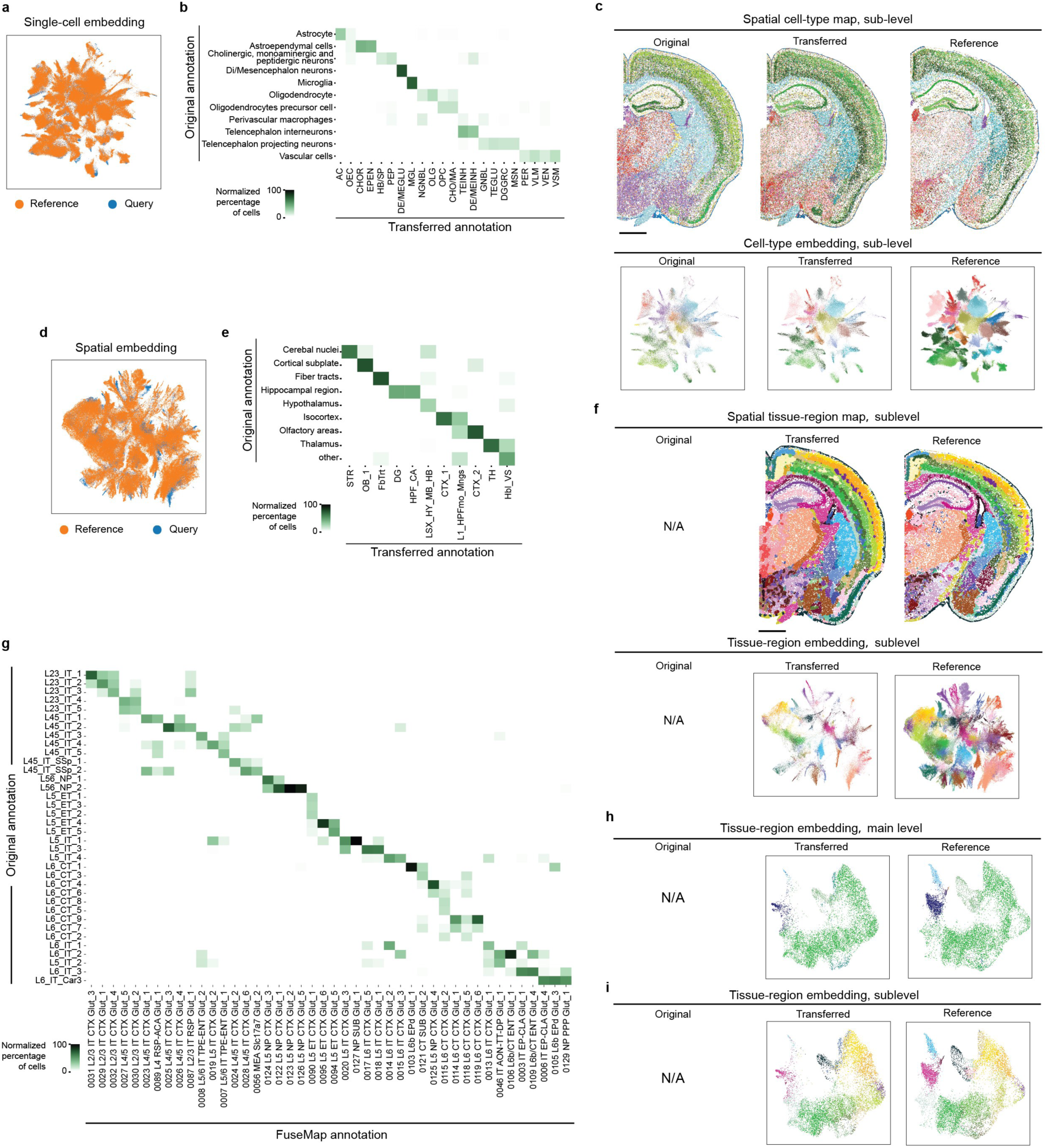
Annotation transfer on query datasets. **a,d**, UMAP of cell-type (**a**) and tissue-region (**d**) embedding colored by the dataset, including one reference sample and one STARmap PLUS query sample^57^. **b**, Percentage of main level original (columns) with transferred (rows) cell types. **c**, Spatial cell-type maps (upper row) and UMAP of cell-type embedding (lower row), colored by original (left) and sublevel transferred (middle) annotations on STARmap PLUS query sample^57^, and sublevel FuseMap (right) annotations on the reference sample. **e**, Percentage of original (columns) with main level transferred (rows) tissue regions. **f**, Spatial tissue-region maps (upper row) and UMAP of cell-type embedding (lower row), colored by sublevel transferred (middle) annotations on STARmap PLUS query sample^57^, and sublevel FuseMap annotations (right) on the reference sample. **g**, Percentage of sublevel original (columns) with transferred (rows) cell types in glutamatergic neurons. **h**,**i**, UMAP of tissue-region embedding at main level (**h**) and sublevel (**i**) annotation on MERFISH query samples^58^ (middle) and the reference sample (right). Scale bar, 1 mm.

**Supplementary Figure 6.**
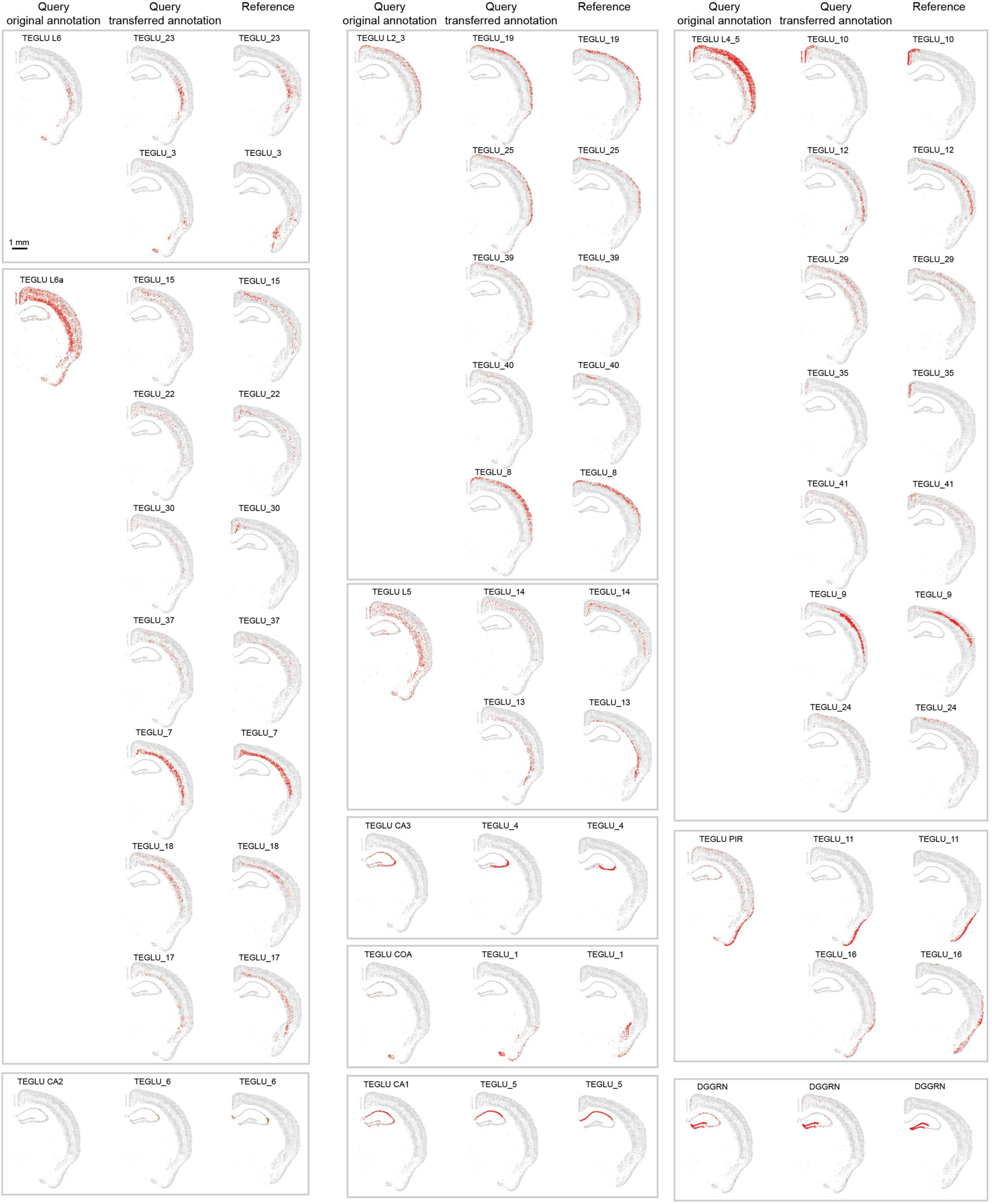
Examples of original and transferred sublevel cell types under TEGLU. Spatial sublevel cell-type maps of original (left) and transferred (middle), annotations on STARmap PLUS query sample^57^, and FuseMap (right) annotations on a reference sample. Scale bar, 1 mm.

